# Multiplexed transcriptomic analyses of the plant embryonic hourglass

**DOI:** 10.1101/2024.04.04.588207

**Authors:** Hao Wu, Ruqiang Zhang, Karl J. Niklas, Michael J. Scanlon

## Abstract

Zoologists have adduced morphological convergence among embryonic stages of closely related taxa, which has been called the phylotypic stage of embryogenesis. Transcriptomic analyses reveal a phylotypic hourglass pattern of gene expression during plant as well as animal embryogenesis, characterized by the accumulation of evolutionarily older and conserved transcripts during mid-embryogenesis, whereas younger less-conserved transcripts predominate at earlier and later embryonic stages. However, widespread comparisons of embryonic gene expression across animal phyla describe an inverse hourglass pattern, where gene expression is correlated during early and late stages but not during mid-development. Here, multiplexed spatial-transcriptomic analyses is used to investigate embryo ontogeny and homology in maize, which has novel, grass-specific morphology. An embryonic-organ genetic network is identified, replete for ancient/conserved genes manifesting a phylotypic hourglass during mid-embryogenesis. Transcriptomic comparisons of grass embryo ontogeny with that of a moss *Physcomitrium patens* identify a conserved, inverse hourglass pattern across plant phyla, as in animals.

The data indicate that the plant phylotypic stage and that of animals is characterized by ancient gene network expression during early histo– and morphogenesis and polarized embryonic development. The data reveal an ancient, convergent mechanism for the evolution of morphological novelty.

---

The embryo is defined as the developmental stage spanning the period between fertilization and the appearance of primary tissues and organs. Post-fertilization, flowering plant embryogenesis consists of three conserved processes: (1) polarized and asymmetric cell divisions to create the embryo proper, (2) patterning of discrete histological tissue-layers, and (3) organogenesis of lateral organs and the formation of apical meristems. Persistent meristematic reservoirs comprising the shoot apical meristem (SAM) and root apical meristem (RAM) enable vascular plants to extend organogenesis beyond embryogenesis. The gene regulatory networks regulating embryogenesis, however, are largely undescribed.

The embryos of the eudicot *Arabidopsis thaliana* and the monocot *Zea mays* ssp. *mays* involve asymmetric cell divisions that establish the embryo proper and the basal, haustorial suspensor that supplies nutrients to the embryo^1–3^ (Fig. 1a). Subsequent histological layering creates an outer epidermal layer surrounding internal tissues, and *Arabidopsis* embryos develop two embryonic leaves (cotyledons) and shoot and root apical meristems, whereupon organogenesis is suspended until seed germination^1, 4^. The grass embryo differs from that of *Arabidopsis*^2, 3, 5–9^ in several important respects. The first lateral organ formed in the maize embryo is the scutellum (Fig. 1b), which at maturity forms a shield-shaped, haustorial organ that absorbs nutrients from the endosperm^7, 10^. The coleoptile then emerges (Fig. 1c) to form a tubular-like sheath that protects the shoot during germination. Unlike *Arabidopsis*, grasses extend embryo organogenesis to develop several foliar leaves contained within the coleoptile before seed quiescence.

**Fig. 1.**
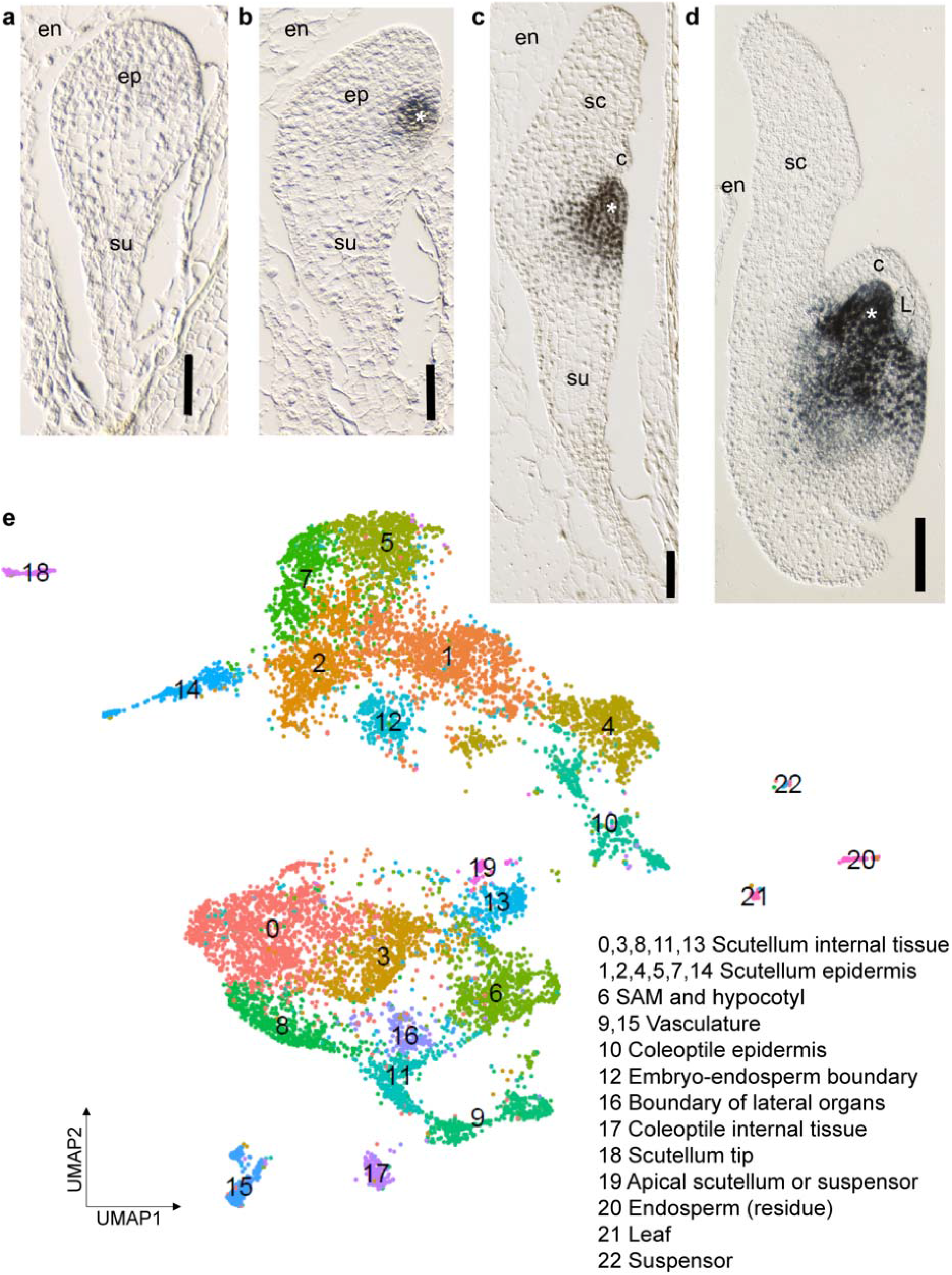
Multiplexed spatial transcriptomics of the maize Stage 1 embryo. Differential interference contrast (DIC) microscopy of maize embryos at the Proembryo stage (**a**), Transition stage (**b**) Coleoptilar stage, (**c**) and Stage 1 (**d**). Immunohistological analyses of the Class I KNOX proteins mark the shoot apical meristem (SAM). ep, embryo proper; en, endosperm; su, suspensor; sc, scutellum; c, coleoptile; L, leaf; *, SAM. Scale bar: 50 μm (**a** and **b**) and 100 μm (**c** and **d**). **e**. Uniform Manifold Approximation and Projection (UMAP) of Stage1 embryo single-cell transcriptomic profiling with cell/tissue types assigned for each cell cluster.

In contrast, the embryos of the moss *Physcomitrium patens* have no apical meristems *sensu stricto* or lateral organs. After fertilization, a transverse zygotic cell division ultimately gives rise to a basal “foot” and a cell mass that develops into the seta and the spore capsule ^11, 12^. The latter consists of a histologically layered sporangium, wherein meiosis results in thousands of haploid spores. Fundamental differences in embryonic growth polarity are also noted between non-vascular and vascular plants, e.g., non-vascular plant embryos develop exoscopically outward from the subtending gametophyte, whereas vascular plant embryos develop endoscopically into the gametophyte^13^. Indeed, embryo morphometric diversity is widespread across the land plants, although comparative transcriptomic analyses of embryogenesis are few^10, 14, 15^.

The diversity of land plant embryo development stands in contrast with that of animals, which appears to converge on to a period of maximum embryo morphological similarity among related vertebrate taxa^16–18^ during “mid-embryogenesis” in contrast to “earlier” and “later” stages that are dissimilar morphologically. The result is an hourglass pattern. The convergent mid-embryo stage has been called the phylotypic stage^19^. Recent transcriptomic analyses of embryogenesis, first performed in fruit flies (*Drosophila*), the zebrafish (*Danio*), and later *Arabidopsis*, have identified a phylotypic stage of gene expression featuring an enhanced accumulation of more ancient and conserved transcripts during mid-embryogenesis as compared to earlier and later ontogenetic stages^20–22^. Plots of transcriptomic gene age and divergence conform to the “hourglass model” first described in animals^16–18^.

However, more widespread comparisons of ontogenetic gene expression across ten diverse animal phyla identify an inverse hourglass pattern, wherein the most prominent transcripts during early and later stages of development are more conserved among phyla as compared to genes expressed during the presumptive “phylotypic stage”^23^.

An important issue in all embryological comparisons across diverse taxa involves temporal homology –– what is meant by early-, mid-, and late-embryogenesis? This issue has added relevance when comparing metazoans and land plants, each of which is adduced to be monophyletic yet morphologically extremely diverse. This concern applies even to broad comparisons among land plant embryos, which manifest what appear to be three, distinct patterns of embryogenesis: the nonvascular bryophytes, the vascular seedless pteridophytes, and the seed plants^13^. Yet, another concern in this field of enquiry is the temporal delay between gene expression and embryological-morphogenetic response, which likely differs among genes within an individual organisms and among phylogenetically diverse taxa. Neither of these concerns has been adequately address, and, as yet a transcriptomic phylotypic analysis is reported for only a single land plant genus (*Arabidopsis*) ^22^. No such study has been reported across the divergent land plant lineages.

Here we present multiplexed analyses of gene expression during maize embryogenesis, a grass that evolved more than 150 million years after the appearance of flowering plants^24^. Single cell sequencing (scRNA-seq), spatial RNA-seq, laser-microdissection RNA-seq (LM RNAseq), and multiplexed RNA-targeting *in situ* hybridizations were utilized to characterize gene expression networks in developing maize embryos at high resolution. An embryonic-organ-initiation genetic network is identified for all lateral organs of the Stage 1 maize embryo, attesting to their collective homology. This embryonic gene network is conserved in *Arabidopsis* and comprises ancient and shared genes expressed in a phylotypic hourglass pattern at mid-embryo stages.

Across-phylum comparisons of embryonic gene expression in the flowering plants maize and *Arabidopsis*, to that of a nonvascular moss, reveals inverse hourglass patterns equivalent to those described in animals. The data indicate that the plant and animal phylotypic stage reveals the ancestral morphometric processes of tissue histogenesis and meristematic growth of embryonic organs. An across plant-animal kingdom model for the evolution of morphological novelty via the innovative expression of newer-evolved, sequence-conserved gene networks is presented.

## Results

### Multiplexed and single-cell profiling of the transcriptomic networks in the Stage 1 maize embryo

A combinatorial sc-RNAseq approach was utilized to identify the gene networks during maize embryogenesis and investigate phylotypic trends and homology of the grass embryo^25, 26^. Stage 1 maize embryos are approximately 1 mm long and contain a prominent, ovately-flattened scutellum, an emerged coleoptile that is beginning to surround the SAM, and the newly-initiated primordium of the first foliar leaf^5^ (Fig. 1d). Stage 1 embryos contain all the lateral organ types of the maize embryonic shoot and thus were an appropriate stage to analyze the organ-specific genetic networks among grass embryonic organs.

10X Genomics Chromium™ was used to profile the single-cell transcriptomes of 11,467 protoplasts digested from over 400 Stage 1 embryos in two replicates (Fig. 1e). Twenty-three cell clusters were identified; median values of 10,594 UMI per cell and 4,030 genes per cell were obtained.

The first challenge encountered in analyses of the sc-RNAseq data was the identification of the tissue/organ-specific origins of each cell-cluster. Prior RNA-seq analyses in maize have focused on seedlings or inflorescences^27^, such that cell-clusters derived from the SAM, embryonic stem, and vasculature^28^ were readily identified, as was the residual starchy endosperm (Fig. 1e and Clusters 6, 15 and 20 respectively). In contrast, relatively few transcriptomic analyses have centered on the grass scutellum or coleoptile, such that the origins of several Stage 1 embryonic cell-clusters were not immediately identifiable. To annotate all the UMAP cell-clusters, a multiplexed transcriptomic approach was used combining the sc-RNAseq data with spatial-transcriptomics and laser-microdissection RNA-sequencing (LM-RNAseq) to determine spatial information for all Stage 1 embryo cell-clusters, as described below (Fig. 2 and Supplementary Fig. 1-2, Supplementary Table 1, 2).

**Fig. 2.**
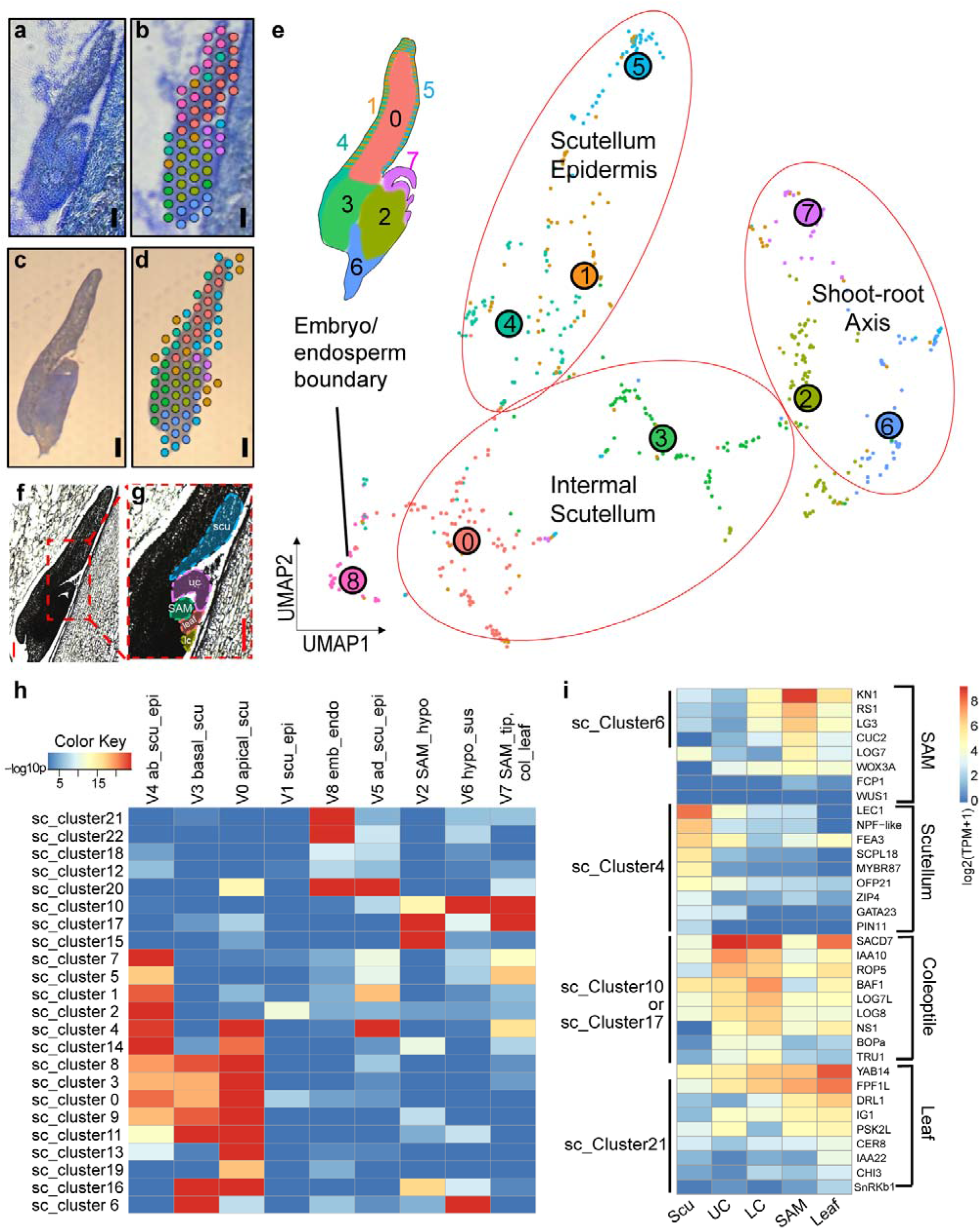
Spatial transcriptomics of maize Stage 1 embryo. Longitudinal cryosections of Stage 1 embryos embedded in OCT (optimal cutting temperature compound; **a-b**) or in paraffin (**c-d)** for 10x Visium^TM^ spatial transcriptomics analysis. **a, c** Toluidine Blue-stained sections for 10x Visium^TM^ spatial transcriptomics assay are overlayed with the10x Visium^TM^ spatial map (**b, d**). The various color-coded circles correspond to the capture spots where spatial cDNA libraries were generated. Scale bar = 100 μm. **e.** Longitudinal sections of paraffin-embedded Stage 1 embryo for use in laser microdissection RNA-seq analysis (LM-RNAseq). **f.** Enlarged view of a Stage 1 embryo showing regions microdissected for LM-RNAseq). Figure labels: scu, scutellum; uc, upper coleoptile; lc, lower coleoptile. **e, f** Scale bar=100 μm. **g**. UMAP of the Visium^TM^ spatial transcriptomic analysis of Stage 1 embryos cell clusters. The spatial, embryonic origins of each cluster are indicated by the color-coded embryo cartoon; numbers correspond to the cluster designations on the UMAP. **h**. Heatmap illustrating the correlation between the single-cell (sc) clusters (y-axis) and Visium^TM^ spatial clusters (x=axis). The significance level is marked by −log_10_p. **i**. Expression of candidate genes identified in microdissected embryonic organs. Cell clusters exhibiting elevated candidate gene expression of the specific markers are indicated; expression values are log_2_ transformed, i.e., log_2_(TPM+1).

To validate the identification of Stage 1 embryo UMAP single cell clusters, eight individual Stage 1 embryo sections were processed using the 10X Genomics Visium^TM^ pipeline for spatial transcriptomics (Fig. 2a-d and Supplementary Table 3). Nine spatially resolved cell-clusters were identified and categorized into four supergroups: (1) shoot-root axis, (2) scutellum epidermis, (3) internal scutellum, and (4) endosperm/embryo boundary (Fig. 2e). Hypergeometric testing was used to analyze the overlap of cluster marker genes between the single-cell and spatial-transcriptomic data, revealing that 14 of the 23 single-cell clusters (0-5, 7-9, 11, 13, 14, 18 and 19) are derived from the scutellum (Fig. 2h). Comparisons of transcript localizations of known marker genes within the UMAP and the spatial-transcriptomic plots indicated that the resolution of Visium^TM^ technology comprises 5-10 embryonic cells. Thus, it was not possible to spatially-distinguish transcripts specifically expressed in the diminutive coleoptile, SAM, or leaf 1. To increase transcriptomic resolution, LM-RNAseq used to microdissect gene expression in the SAM, adaxial scutellum, upper/lower coleoptile, and leaf primordium of Stage 1 embryos (Fig. 2f-g). Principal component analyses (PCA) showed that the transcriptomic profiles of these five, maize embryonic organ/tissues are distinct (Supplementary Fig. 2a). In particular, 9,913 genes were identified as differentially-expressed in at least one embryonic structure, and that 89, 1058, 246, 84, and 53 genes were preferentially expressed in these same five embryonic organ/tissues (Supplementary Fig. 2b-f and Supplementary Table 4-6). Thus, the multiplexed transcriptomic approach enabled the identification of the tissue/organ origin of each cell-cluster in the Stage 1 maize embryonic UMAP (Fig. 1e, 2i).

### Spatiotemporal analyses of embryo ontogeny and homology via multiplexed RNA-targeting

Homologous organs are thought to employ shared gene networks regulating developmental patterning^25, 29^. With an emphasis on regulatory genes differentially expressed in our transcriptomic analyses of Stage 1 maize embryos, a total100 transcripts were targeted for spatiotemporal analyses of multiplexed, *in-situ* hybridization of transcript localization during embryo ontogeny, using the 10X Xenium^TM^ protocols (Supplementary Table 7). Samples included longitudinal sections of (i) transition-staged embryos just forming a SAM and scutellum (Fig. 1b), (ii) coleoptilar-staged embryos (Fig. 1c), and (iii) Stage 1 embryos initiating the first foliar leaf (Fig. 1d). Correlations of the sc-RNAseq data, spatial-transcriptomic plots, and Xenium^TM^ *in situ* RNA-targeting data were examined via comparisons of 16 genes preferentially expressed in one-or-two single-cell UMAP clusters (Supplementary Fig. 1). In all cases, the *in situ* hybridization patterns were consistent with the spatial expression data.

In light of the comparative consistencies of the multiplexed *in situ* hybridization data, the dynamic co-localization patterns of some key regulators of maize shoot development during serial stages in grass embryo ontogeny were examined. Consistent with prior studies^30, 31^, the spatial-temporal expression of the indeterminacy marker *KNOTTED1* (*KN1*; Fig. 3a-d) was first detected during SAM initiation at the maize Transition stage (Fig. 1b). At later stages, *KN1* was observed to accumulate in the SAM and subtending embryonic stem, but was downregulated in lateral organs^30, 31^. *NARROW SHEATH1* (*NS1*) and *WUSCHEL-RELATED HOMEOBOX 3A* (*WOX3A*), which encode homeodomain transcription factors required for mediolateral expansion of maize lateral organs^29^, were co-expressed at the edges of the developing coleoptile and initiating leaf primordia as described previously^7, 25, 32, 33^ (Fig. 3b-c, f-g). Undetected in prior studies of grass embryo ontogeny, Fig. 3a reveals that *NS1* transcripts accumulate in the initiating scutellum at the Transition stage. Moreover, transcripts of *CUP-SHAPED COTYLEDON2* (*CUC2*), which mark developmental boundaries throughout maize ontogeny^34^, were detected between the expression domains of *KN1* and *NS1* during scutellum initiation (Fig. 3a), and in the boundaries between the SAM and later-staged embryonic lateral organs (Fig. 3b-c,e). *DROOPING LEAF1* (*DRL1*), which regulates leaf architecture and is expressed in incipient and developing foliar leaf primordia^35^, was not detected in the coleoptile or scutellum at any embryo stage examined (Fig. 3c,h). These combinatorial *in situ* RNA-targeting analyses enabled the identification of previously undescribed combinatorial patterns of dynamic, gene expression during maize embryo development.

**Fig. 3.**
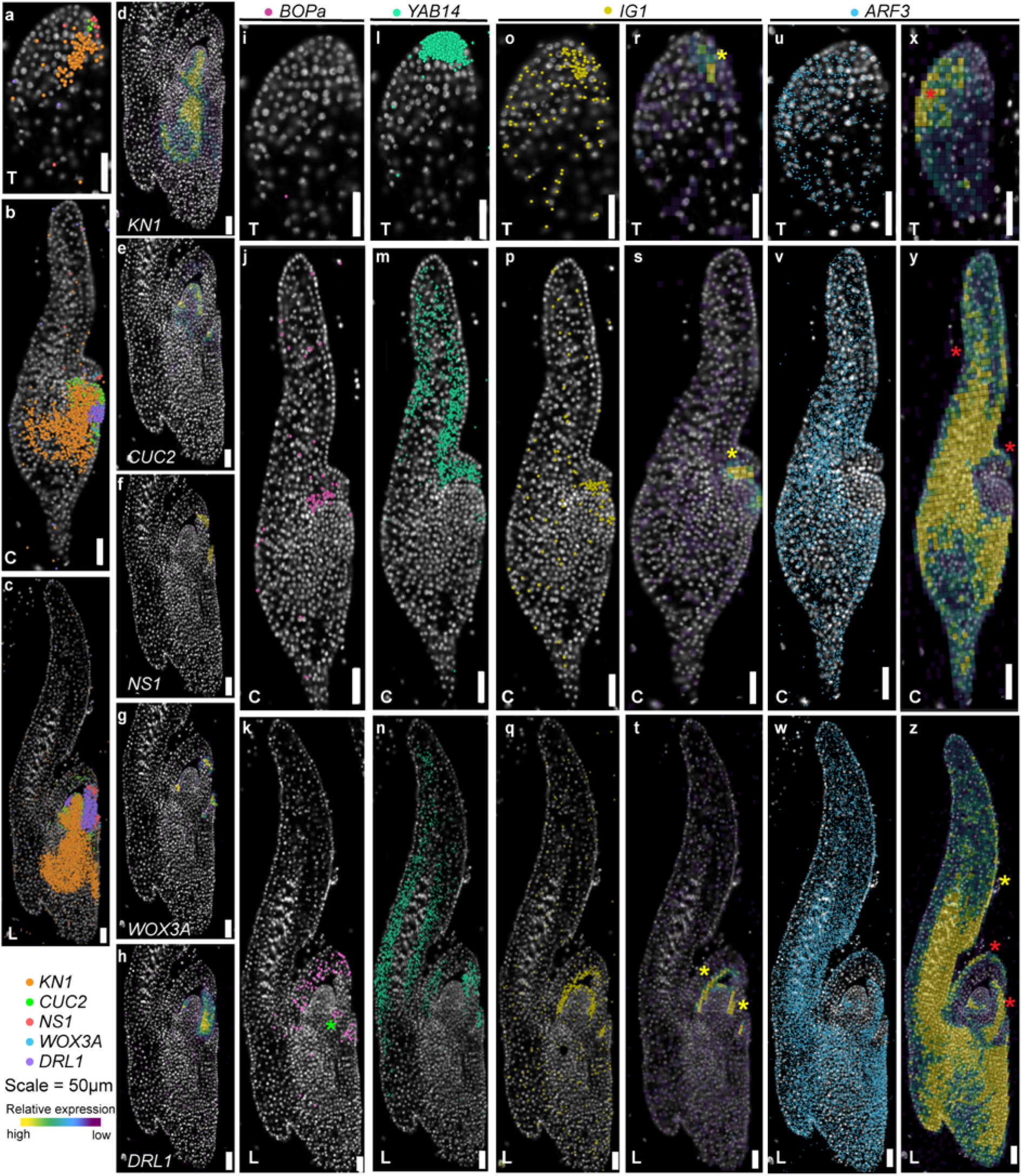
RNA-targeting analyses of Transition-staged, Coleoptilar-staged, and Stage 1 embryos. **a-c**. Signal point map of *KN1, CUC2, NS1, WOX3A* and *DRL1* at the Transition stage (**a**), Coleoptilar stage (**b**) and Stage 1 (**c**). **d-h.** Expression density map of *KN1* (**d**)*, CUC2* (**e**)*, NS1* (**f**)*, WOX3A* (**g**) and *DRL1* (**h**) at Stage 1. **i-k, l-n, o-q** and **u-w**. Signal point map of *BOPa* (**i-k**), *YAB14* (**l-n**), *IG1* (**o-q**), and *ARF3* (**u-w**) at the Transition stage, Coleoptilar stage and Stage 1. **r-t** and **x-z**. Expression density map of *IG1* (**r-t**) and *ARF3* (**x-z**) at the corresponding stages. Figure labels: T, Transition stage; C, Coleoptilar stage; L: Stage 1. The point colors are arbitrarily determined. Yellow asterisks mark the adaxial expression of *IG1* and *ARF3*, and red asterisks mark the abaxial expression of *ARF3*. The green asterisk in panel **k** marks the expression maximum of *BOPa* at the emerging leaf sheath. The yellow asterisks in panel **r-t** and **z** mark the expression maxima of *IG1* at the adaxial side of the scutellum (**r**), the coleoptile (**s**), the coleoptile and the leaf (**t**), as well as the expression maxima of *ARF3* at the adaxial side of the scutellum (**z**), respectively. The red asterisks in panel **x-z** mark the abaxial expression of *ARF3* at the scutellum (**x, y**), the coleoptile (**y, z**) and the leaf (**z**). Scale bar for all panels = 50 μm.

Other candidate genes examined using Xenium^TM^ displayed patterns of transcript accumulation that offer insights into the developmental homologies of maize embryonic lateral organs. For example, *BLADE-ON-PETIOLEa* (*BOPa*) accumulated throughout the developing coleoptile and in the proximal region of the initiating leaf at Stage 1 (Fig. 3i-k), consistent with the proposed function of *BOP* homologs as leaf proximal-identity genes^36^. Notably, *BOPa* was undetected throughout scutellum ontogeny, in a manner similar to the accumulation patterns noted for the *BOPa* paralog *TASSELS REPLACE UPPER EARS1* (*TRU1*) (Supplementary Fig. 3). Transcripts of the *DRL1-*paralog *YABBY14* (*YAB14*) densely accumulated in the apical tip of the scutellum at the transition stage and in the elongating scutellum and newly emerged coleoptile at the coleoptilar stage. During Stage 1, a new *YAB14* expression maximum was organized at the primordial leaf tip (Fig. 3l-n). Moreover, the pleiotropic gene *INDETERMINATE GAMETOPHYTE1* (*IG1*), reported to function during leaf polarity and patterning of maize leaves^37, 38^, accumulated on the dorsal (i.e. adaxial) surfaces of initiating leaf primordia during Stage 1, and also in the adaxial layers of the emerging coleoptile and in the upper-adaxial region of the transition-staged scutellum (Fig. 3o-t). No significant *IG1* expression was observed during later stages of embryonic scutellum development (Fig. 3p-q, s-t). Another leaf polarity regulator, *AUXIN RESPONSE FACTOR 3* (*ARF3*)^39^, was expressed on the abaxial region of the transition-staged embryo during scutellum initiation, and in the abaxial domains of the newly-emerged coleoptile and leaf during later embryo stages (Fig. 3u-z). Interestingly, the expression of *ARF3* in the scutellum moved from the ventral (i.e., abaxial) region in the transition stage to adaxial region at Stage 1 (Fig. 3z).

### The initiation of maize embryonic lateral organs utilizes a shared genetic network

By Stage 1, the three grass embryonic lateral organs initiate morphological divergence and continue to do so before seed quiescence^5^. Therefore, the determination of organ homology and of a transcriptomic phylotypic stage in maize requires the examination of genetic networks throughout embryo ontogeny^26^. Toward this end, the LM-RNAseq data from Stage 1 embryonic organs were combined with the LM-RNAseq data from serial stages of maize embryonic development^28^ to generate a more comprehensive sampling throughout embryogenesis. Specifically, these prior analyses generated transcriptomic data precisely from the newly initiated embryonic organ and the adjacent SAM, which were serially sampled at the transition stage (scutellum +SAM), the coleoptilar stage (coleoptile +SAM) and Stage 1 (leaf 1 +SAM). Fig. 2i shows that the expression levels of several leaf developmental genes were highest in Stage 1 leaves (L) and steadily decreased in the lower coleoptile (LC), upper coleoptile (UC), and scutellum (S). Notably, the expression of these genes during the transition stage and the coleoptilar stage was similar to that of leaf 1 during Stage 1 (Supplementary Table 4)^28^. These data provide evidence that the gene networks involved in foliar leaf initiation at Stage 1 are also associated with the initiation of the scutellum and coleoptile at successively earlier stages in embryo ontogeny.

To further investigate this hypothesis, a weighted gene co-expression network analysis (WGCNA)^40, 41^ was performed on the Stage 1 embryonic LM-RNAseq data to identify co-expression modules correlated with the expression pattern L > LC > UC > S. A total of 5,797 gene transcripts with high expression variability were selected. With the exception of one non-clustered transcript module (MEgrey module), these were grouped into twelve co-expression modules (ME1 through ME12). Two co-expression modules (ME2 and ME3) comprising 963 genes emerged as correlated with the L > LC > UC > S pattern. (Supplementary Fig. 4a and Supplementary Table 8). Intersection of these 963 genes with transcripts identified by LM-RNAseq in the initiating scutellum, coleoptile and leaf 1^28^ (excluding putative SAM-specific genes) identified 130 genes comprising the maize embryonic-organ-initiation gene network (Fig. 4a and Supplementary Fig. 4b and Supplementary Table 9). Expression of this gene network was significantly higher during initiation of these embryonic organs as compared to later morphometric stages (Fig. 4b, Supplementary Fig. 4b-e).

**Fig. 4.**
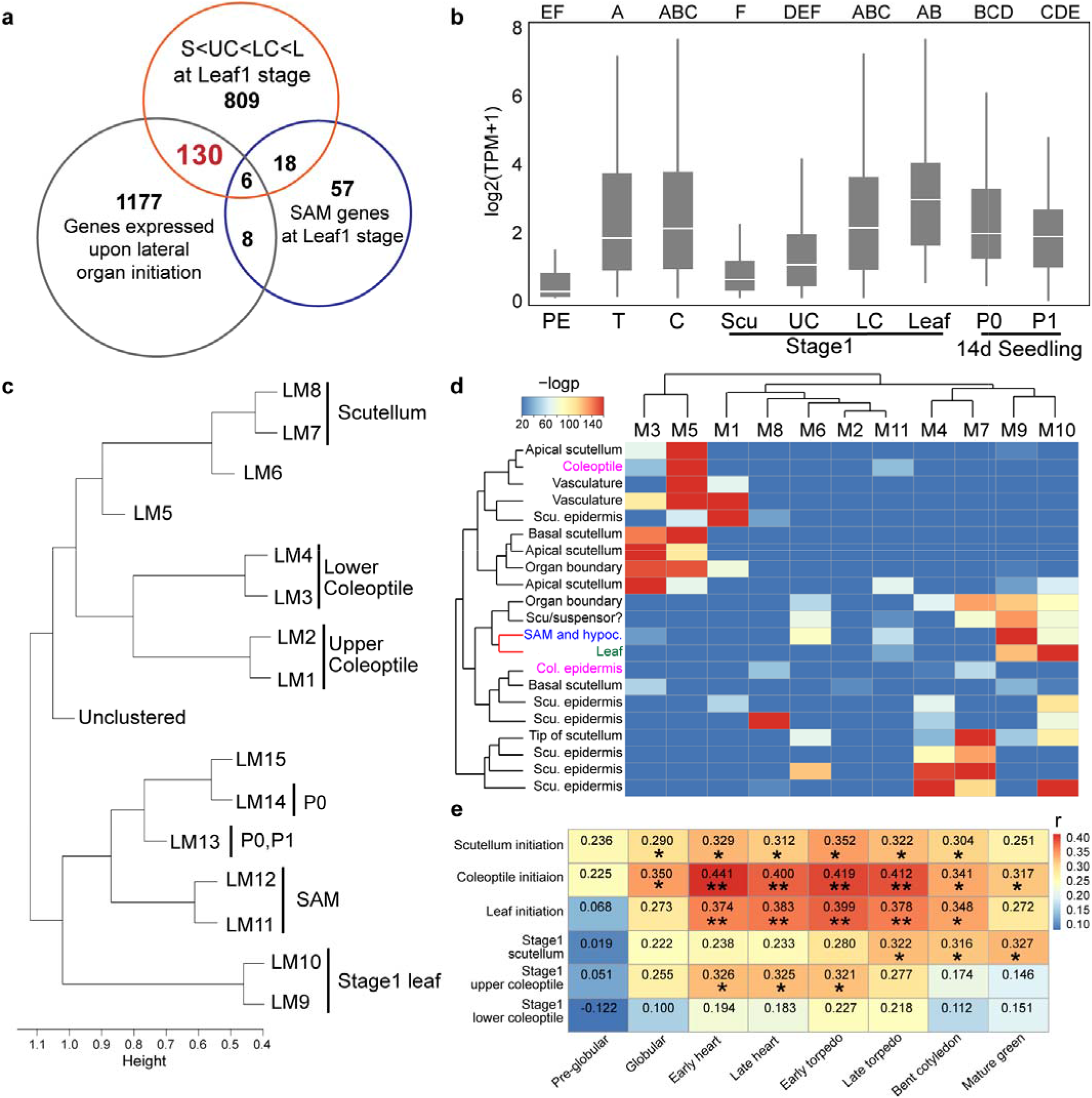
Dynamics of the maize embryo organ initiation network. **a**. A Venn diagram of embryonic transcriptomics identifies 130 genes in the embryo-organ-initiation-network. **b**. Normalized expression of these 130 genes at the Proembryo stage (PE), Transition stage (T), Coleoptilar stage (C), Stage 1 scutellum (Scu), Stage 1 upper coleoptile (UC), Stage 1 lower coleoptile (LC), Stage 1 leaf (L), as well as P0 and P1 of the 14 day-after-germination seedlings. The letters above the boxplot represent significance levels (non-overlapping letters indicate significant differences at p<0.05) using the Tukey-Kramer HSD test. **c.** WGCNA tree graph of the transcriptomic relationships among organ-specific co-expression modules. The height (y-axis) scales the length of organ-specific branches, indicating the transcriptomic distances between modules. **d**. Heatmap illustrating correlations between single-cell (sc) clusters (y-axis) and hdWGCNA modules (x=axis). The significance levels are marked by −log_10_p. **e**. Heatmap illustrating the correlation of the initiation network expression between Maize (y-axis) and Arabidopsis (x-axis) across the embryogenesis. Heatmap colors are determined by correlation coefficient (r) via Pearson correlation test with significance level p<0.05 (*) or p<0.01 (**). Scu, Scutellum; Col, coleoptile; hypoc, hypocotyl.

In light of the controversy regarding the homology of the grass cotyledon (i.e., whether the coleoptile is a foliar leaf^8^, the entire cotyledon^9^, or the proximal component of a bipartite, single, grass cotyledon^3, 33^), a second WGNCA analysis was performed on the LM-RNAseq data obtained from the Stage 1 embryonic shoot apex. In addition, published LM-RNAseq data^42^ of the Plastochron 0 (P0) and P1 leaves, from 14-day after planting maize seedlings, were integrated as a reference for later leaf ontogeny. Batch effects were removed (Supplementary Fig. 5a,b), and the merged dataset was filtered to select genes with variability in transcript accumulation levels across embryonic organs. In the analysis, 5,128 out of 5,137 genes were selected and clustered into 15 co-expression modules (LM1, LM2, …, LM15). The remaining 9 transcripts were resolved into an unclustered module (Supplementary Table 10). Transcriptomic relationships among the co-expression modules are shown in a tree graph (Fig. 4c), in which the y-axis (height) scales the length of organ-specific branches, indicating transcriptomic distances between these modules (Supplementary Fig. 5a,b). Pearson correlation tests between module eigengenes and organ-specific transcriptomes identified associations between specific co-expression modules and the corresponding embryonic organs (Fig. 4d and Supplementary Fig. 5c). Two major branches comprise (1) the Stage 1 leaf, SAM, P0 and P1, and (2) the coleoptile and the scutellum.

To further investigate the transcriptomic relationships among all Stage 1 maize embryonic lateral organs at single-cell resolution, high-dimensional WGCNA (hdWGCNA^43^) was performed on the scRNAseq data. A total of 5,026 genes transcripts were clustered into 11 co-expression networks (M1, M2, …, M11; Supplementary Table 11). A hypergeometric test was used to analyze the overlap between the putative single-cell cluster marker-gene transcripts and those highly correlated with corresponding co-expression modules. The data indicate that the SAM and the leaf cluster marker-genes significantly overlap with the closely related modules M9 and M10, respectively. In contrast, marker genes from the scutellum and the coleoptile cluster overlap with multiple co-expression modules (M5, M7 and M8; Fig. 4d). The single-cell cluster tree shows that the coleoptile together with the scutellum and the leaf is next to the SAM, consistent with the LM-RNAseq WGCNA data. (Fig. 4c).

The WGCNA clustering results linking the scutellum and coleoptile were supported by genetic analysis. *LONELY GUY7* (*LOG7*), which encodes a cytokinin-activating required for maize SAM maintenance^42^, was expressed at the SAM tip in Stage 1 embryos (Supplementary Fig. 6a-c). Null mutations in *log7* resulted in SAM termination^42^. Histological examination of *log7* mutant Stage 1 embryos revealed the absence of a detectable SAM or foliar leaf, although the scutellum and coleoptile were morphologically normal (Supplementary Fig. 6d-h). These data expose morphogenetic differences between leaves, and the scutellum and coleoptile.

### The embryonic-organ-initiation genetic network stage is conserved in Arabidopsis embryos

To investigate the transcriptomic similarities of embryonic lateral organs of maize and those of the eudicot model Arabidopsis, comparative genomic toolkits [GRAMENE (https://www.gramene.org/), TAIR (https://www.arabidopsis.org/), BLAST tools (https://blast.ncbi.nlm.nih.gov/Blast.cgi)] were used to select *Arabidopsis* orthologs of the 130 transcripts in the maize embryonic-organ-initiation network, and to investigate the gene expression atlas of *Arabidopsis* embryo development^44^. Embryonic nomenclature differs between eudicots and grasses^1^. In *Arabidopsis*, histological layering and SAM initiation occur during the globular stage, cotyledon initiation and emergence defines the heart stages, whereas the cotyledons elongate during the torpedo and bent embryo stages.

Out of 130 genes in the maize embryonic-organ-initiation network, forty-nine had corresponding *Arabidopsis* orthologs expressed during embryo development, commencing during cotyledon initiation at the early heart stage and persisting until the late cotyledon stage (Supplementary Table 12). Notably, the stage-specific accumulation patterns of these homologous maize transcripts in the emerging scutellum (Transition stage), coleoptile (Coleoptilar stage), and foliar leaf (Stage 1) were significantly correlated with the reported expression of the corresponding *Arabidopsis* orthologs during specific stages in cotyledon initiation and development in this eudicot embryo^44^ (Fig. 4e, Supplementary Fig. 7). In addition, transcripts accumulating in the expanding Stage 1 scutellum correlated with expression of *Arabidopsis* orthologs during the later stages of cotyledon growth (i.e. late-torpedo stage^44^). Likewise, maize transcripts accumulating in the elongating Stage 1 coleoptile were significantly correlated with expression of orthologous *Arabidopsis* genes during earlier stages of cotyledon expansion (i.e., the early heart stage^44^; Fig. 4e). These data were interpreted to indicate that despite the morphological dissimilarities of these grass and eudicot “mid-staged” embryos, the development of the maize and *Arabidopsis* embryonic lateral organs share a homologous regulatory network.

### The embryonic organ initiation network comprises a phylotypic stage

The data reported in this study support the hypothesis that the maize embryonic-organ-initiation network is conserved in *Arabidopsis thaliana* and correlates with the transcriptomic signatures of the eudicot heart and torpedo stage embryos. Prior studies report that during mid-embryogenesis, old and conserved genes are more highly expressed than young and divergent genes, marking the transcriptomic phylotypic stage of *Arabidopsis* embryogenesis^22^. To investigate if the maize embryonic-organ-initiation network also comprises a transcriptomic phylotypic period in grass embryogenesis, the age and divergence of the genes in the maize embryonic-organ-initiation network were studied.

To evaluate gene age, a phylostratigraphic analyses obtained 13 phylostrata (PS), PS1-PS13 (Fig. 5a and Supplementary Table 13) using statistical genetic methods to identify homologous founder genes across lineages (including 350 species) and date their phylogenetic origin (Supplementary Table 14)^45^. Significantly, 102 of the 130 transcripts in this phylotypic maize embryonic-organ-initiation-network appear in one of three phylostrata, i.e., phylostratum PS1 (29; unicellular eukaryotes and prokaryotes), PS2 (36; all eukaryotes), or PS4 (37; Bryophyta). Less than 10 percent of this maize embryonic network comprises newer transcripts (from PS5-PS13) that appear to have arisen later in plant evolution (Supplementary Figs. 8a-c). Notably, the majority of genes in this network appear within the Bryophyte phylostratum PS4, which marks the first appearance of the plant embryo^24^. In contrast, just 12 embryonic transcripts (9.23%) are identified in the algal phylostratum (PS3), which consists of plants lacking embryos. Gene expression numbers during the non-phylotypic Proembryo stage are dominated by later evolved genes from PS9-PS11 (i.e., the Poaceae or grasses, the panicoid grasses, and the genus *Zea*; Fig. 5a), whereas transcripts from Stage 2 embryos and seedling leaves are enriched for newly evolved genes from PS12 and PS13 and the genus *Zea* (Supplementary Table 15).

**Fig. 5.**
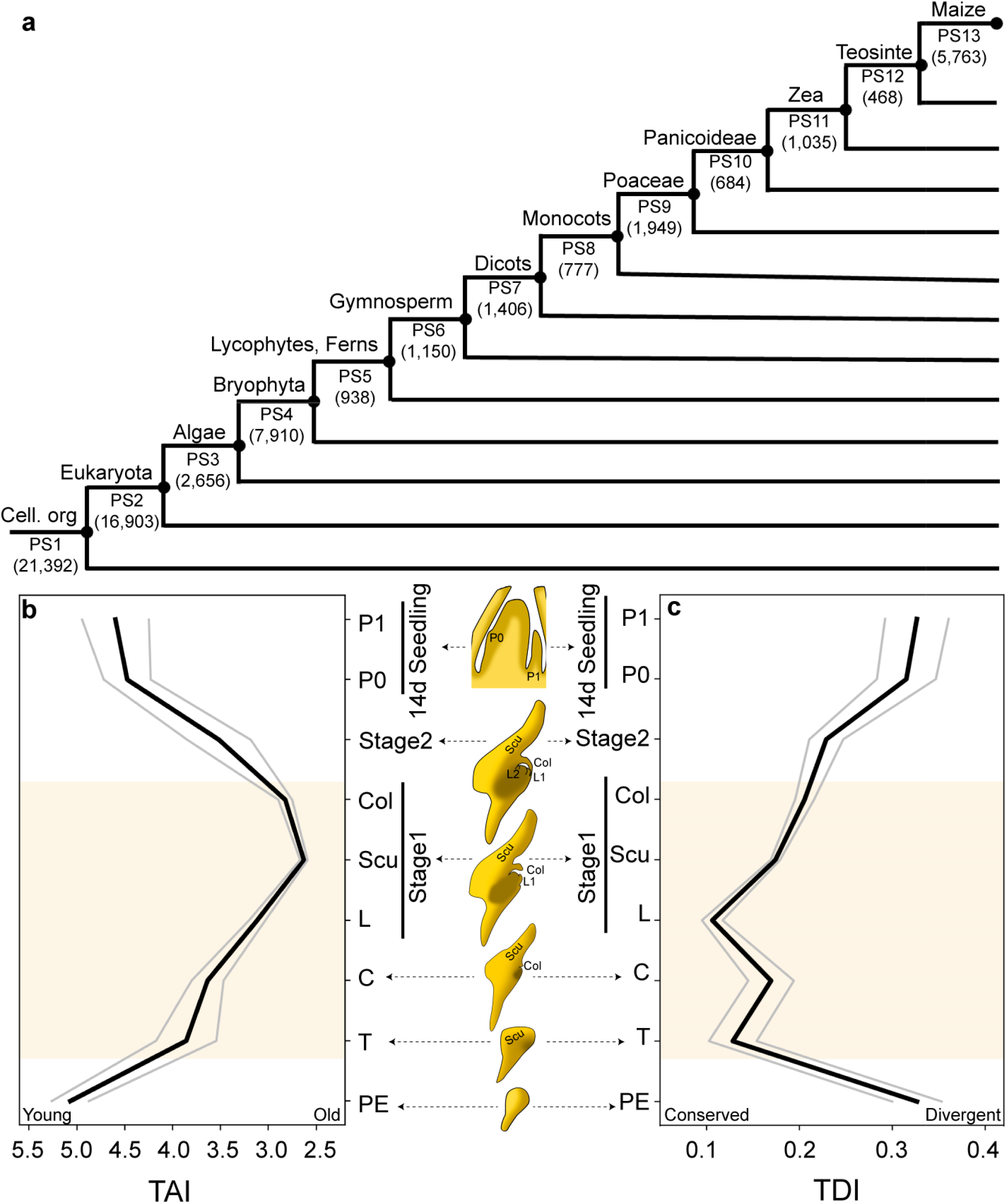
Phylotypic stage analysis of the maize embryogenesis. **a**. Phylostratigraphic map of maize. All maize protein-coding transcripts were assigned to thirteen phylostrata (PS1-PS13). Parentheses designate the number of genes per phylostratum; taxonomic groupings are marked above the corresponding PS. **b-c**. The transcriptome age index (TAI; **b**) and transcriptome divergence index profiles (TDI; **c**) during maize embryogenesis and formation of seedling leaves after germination. Figure labels: PE, proembryo stage; T, transition stage; C, coleoptilar stage; L, Stage 1 leaf; Scu, Stage 1 scutellum; Col, Stage 1 coleoptile; P0 and P1, Plastochron stages of the 15th and 14th leaves in the 14-day after planted seedlings. The gray lines represent the standard deviation estimated by bootstrap analysis as described^22^. The overall patterns of TAI and TDI profiles are significant, as measured by the flat line test (*P*_TAI_=3.56×10^-6^, *P*_TDI_=4.80×10^-4^), reductive hourglass test (*P*_TAI_=1.49×10^-4^, *P*_TDI_=1.53×10^-3^), and reductive early conservation test (*P*_TAI_=1, *P*_TDI_=0.998).

The transcriptome age index (TAI) measures gene age adjusted by gene expression level for each embryogenic stage (Supplementary Table 16). As shown in Fig. 5b and Supplementary Fig. 8c, the TAI displays a “classic” hourglass pattern featuring a significant decrease beginning at the proembryo stage that reaches a minimum value during Stage 1, followed by an increased TAI at Stage 2 that continues in the newly initiated leaves of later, germinated maize seedlings. The transcriptome divergence index (TDI) measures the average sequence divergence of genes, weighted by transcript accumulation level, at different developmental stages (Supplementary Table 17). Consistent with the TAI results, the TDI also showed an hourglass pattern that commences at the Proembryo stage and extends to Stage 1, followed by a notable increase at Stage 2 and in germinated seedling leaf primordia (Fig. 5c and Supplementary Fig. 8d-e). Thus, measures of both transcriptional gene age and gene sequence divergence reveal a mid-embryogenic, phylotypic “hourglass” pattern during maize embryogenesis. The data also reveal that the phylotypic-staged gene networks found in maize and *Arabidopsis* are homologous (Fig. 4e), as is predicted for two seed plants from the same morphological phylotype^13, 23^.

### Transcriptomic phylotypic comparisons across plant phyla reveal an inverse hourglass pattern

A comprehensive transcriptomic survey of embryonic/sporophytic development in the non-vascular plant *Physcomitrium patens* is available for use in across-phylum analyses of phylotypic gene expression in plants^46^. Ortiz-Ramirez et al., (2016)^46^ sampled gene expression in the zygote and young embryo housed within the gametophytic archegonium, and from four subsequent ontogenetic increments in moss embryo development (labeled S1-3, and SM). As described in Janzen (1929)^11^ and modeled in Fig. 6a, key morphogenic events in *P. patens* sporophyte development include the establishment of a multicellular embryo during S1 [5-6 days after fertilization (AF)] and subsequent proliferative growth of the seta and spore capsule during both the S2 (9-11 days AF) and S3 stages (18-20 days AF) (Fig. 6a). Formation of the sporangial epidermal layers occurs during S3, followed by meiosis in the mature sporangium during the SM stage (28-30 days AF). It is important to note that the archegonium samples consist of primarily gametophytic, not embryonic/sporophytic, tissues^46^.

**Fig. 6.**
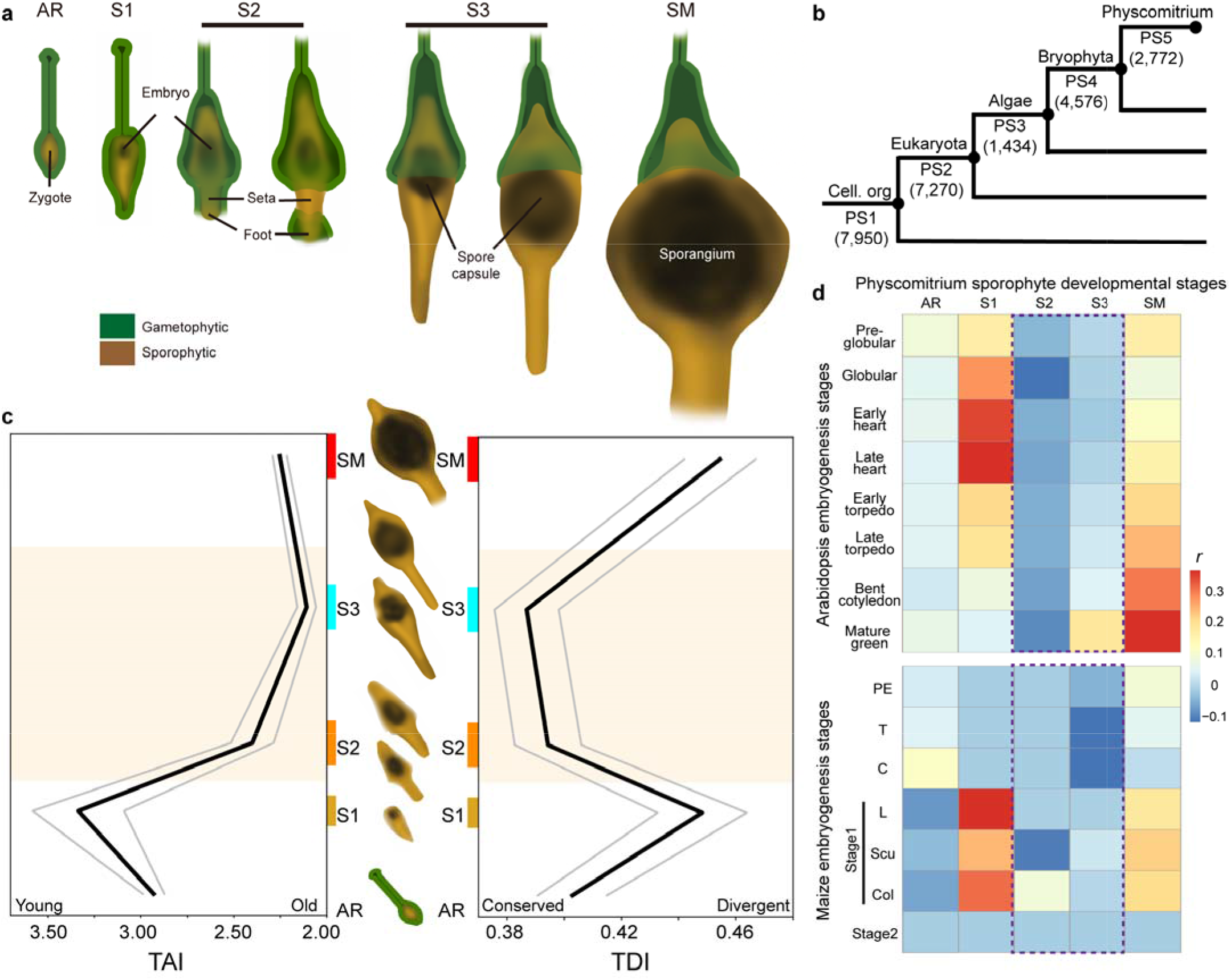
Across-phylum phylotypic analyses of *Physcomitrium patens.* **a.** Development of embryo/sporophyte in the moss *P. patens*. **b**. Phylostratigraphic map of *P. patens*. All *P. patens* protein-encoding transcripts were assigned to one of five phylostrata (PS1-PS5). Parentheses refer to the number of genes per phylostratum; taxonomic groupings are noted. **c**. The transcriptome age index (TAI) and transcriptome divergence index (TDI) profiles of *P. patens* archegonia; sporophyte stages S1, S2, S3 and SM. Colored bars illustrate the duration and timing of different S stages along the 30 day time scale. **d**. Comparative expression levels of stage-specific embryonic genes between *Arabidopsis* and *P. patens,* and maize and *P. patens*. Figure labels: (in moss) AR, archegonium; S1, sporophyte 1 stage; S2, sporophyte 2 stage; S3, sporophyte 3 stage; SM, mature sporophyte stage; (in maize) PE, proembryo stage; T, transition stage; C, coleoptilar stage; L, Stage 1 leaf; Scu, Stage 1 scutellum; Col, Stage 1 coleoptile; S2, Stage 2. The gray lines represent the standard deviation estimated by bootstrap analysis as described^22^. The overall patterns of TAI and TDI profiles are significant, as measured by the flat line test (*P*_TAI_=0.0131, *P*_TDI_=4.73×10^-3^), reductive hourglass test (*P*_TAI_=0.0332, *P*_TDI_=0.0249), and reductive early conservation test (*P*_TAI_=0.998, *P*_TDI_=0.973). Images in (**a)** and (**c**) are reproduced from Ortiz-Ramirez et al., (2016)^46^.

Five phylostrata were used to determine the ages of genes enriched in *P. patens* embryos, whereas their sequence divergence (K_a_/K_s_) ratios were calculated by comparing gene homologs in *Physcomitrellopsis africana* (Fig. 6b and Supplementary Table 18-20). TAI and TDI were rendered for transcripts enriched during each of the five developmental increments: archegonia, and sporophyte stages S1, S2, S3, and SM. The data indicate an increasing TAI and TDI pattern from the archegonium stage to S1, whereas older embryonic transcripts increase markedly during S2, reaching a phylotypic peak at S3, followed by increasing TAI and TDI at SM reflecting the use of relatively younger genes in the mature sporophyte (Fig. 6c). Thus, embryos of this moss also manifest a phylotypic hourglass pattern of gene expression at mid-embryogenesis.

To test if embryos of *P. patens* utilize a similar phylotypic gene network as the seed plants *Arabidopsis* and maize, correlations of embryonic gene expression were made across phyla. The data reveal an inverse hourglass pattern (Fig. 6d), with little correlation between mid-embryonic phylotypic-staged transcripts of the moss (S2-S3) and genes enriched during the phylotypic periods of maize or *Arabidopsis*. In contrast, early (S1) and later-staged (SM) embryonic networks in the moss show increased correlations with genes expressed in the seed plant embryos. Thus, as described in animal embryos, across-phylum comparisons of plant embryonic gene expression yield an inverse hourglass pattern characterized by the expression of distinct, phylotypic gene networks.

## Discussion

A multiplexed transcriptomic approach was applied to investigate embryogenesis in a monocot grass plant, which has evolved novel embryonic structures with previously undefined gene expression networks. Spatial transcriptomic technologies complement the ambiguities encountered in cell-cluster identification that can otherwise complicate scRNA-seq analyses, whereas the precision of LM-RNAseq can delineate the tissue/organ-specific transcriptomes of embryonic organs that are unresolved using spatial transcriptomics alone (Fig. 2). Moreover, high-throughput, multiplexed *in situ* RNA-targeting technologies enable localizations of multiple transcripts simultaneously and combinatorically, throughout embryo ontogeny. The RNA-targeting analyses presented in this study support the expression of a gene network that functions during the initiation of the grass scutellum, coleoptile, and foliar leaf (Fig. 3). Thus, the combinatorial application of multivariate transcriptomic approaches enables a high-throughput and spatially-informed transcriptomic profile of grass embryo development at cellular resolution (Fig. 1).

Although questions about the evolution and homology of the cotyledon remain unresolved, genetic and morphological models indicate that the cotyledon evolved via the modification of a first embryonic leaf^1, 4, 47^. In support of these models, our transcriptomic comparisons of maize embryonic lateral organs identify a conserved embryonic-organ-initiation network (Fig. 4a), consistent with the proposition that the scutellum, coleoptile, and leaf 1 are indeed homologous organs. Likewise, our co-expression analyses, RNA-targeting assays, and genetic analyses of meristem mutants collectively support the hypothesis that the cotyledon is a bipartite organ comprising an apical, haustorial scutellum fused to a basal, emergent and sheathing coleoptile^3,33^. The scutellum and coleoptile cluster together in WGCNA analyses (Fig. 4c) and distant from foliar leaves, reflecting the expression of genetic networks that are more closely-shared within these first-elaborated embryonic organs as compared to leaves. Genetic analyses also reveal that *Zmlog7* mutants fail to make a SAM or leaves, although the scutellum and coleoptile are unaffected, likewise consistent with morphogenetic differences between leaves and both the scutellum and coleoptile (Supplementary Fig. 6). Similar phenotypes are reported for null mutations of the meristem maintenance gene *KNOTTED1,* wherein the SAM and leaf are absent, but the coleoptile and scutellum are unperturbed by loss of leaf and meristem integrity^30^. Furthermore, expression of the proximal, sheath marker genes *BOPa* and *TRU1*^34,38, 48^ is seen in the Stage 1 coleoptile but not in the scutellum (Fig. 3i-k and Supplementary Fig. 3), in support of models where the coleoptile is purported to be homologous to the proximal, leaf sheath and the scutellum comprises the highly modified distal component of a fused, bipartite, grass cotyledon^3, 33^.

Further, our multiplexed transcriptomic analyses of maize embryogenesis identify a genetic network expressed during initiation of embryonic organs that describes a phylotypic period (Fig 5b-c). As previously reported in the eudicot *Arabidopsis* and in the embryos of fruit flies (*Drosophila melanogaster*) and zebrafish (*Danio rerio*)^20–22^, the transcriptomic phylotypic period is defined by the preferential expression of evolutionarily ancient and sequence-conserved genes. We note that the transcriptomic phylotypic period in maize embryogenesis commences during the late proembryo stage and forms an extended hourglass shape corresponding to successive initiations of the scutellum (at transition stage), coleoptile (at coleoptilar stage), and leaf 1 (at Stage 1). Significant decreases in both transcript age and conservation occur during later stages of maize embryogenesis (Stage 2), and upon the initiation of seedling leaves after germination (Fig. 5b-c).

As in the seed plants maize and *Arabidopsis*, embryos of the non-vascular moss *Physcomitrium patens* display transcriptomic phylotypic hourglass patterns that peak mid-embryonically (Fig. 6c). In this context, it is worth noting that the phylotypic-stage gene network in *P. patens* consists of transcripts that are markedly distinct from the gene networks expressed during phylotypic mid-embryogenesis in *Arabidopsis* and maize. Indeed, across-phylum transcriptomic comparisons display an inverse hourglass pattern, where the least homologous embryonic gene expression networks occur at mid-embryonic stages, whereas early and later stages of moss sporophyte/embryo development show increased co-expression with maize and *Arabidopsis* embryonic networks (Fig. 6d). These findings are consistent with across phylum phylotypic comparisons in animals^23^, and likewise support predictions that non-vascular plants and seed plants comprise distinct, embryonic phylotypes^13^.

Historically, morphological and transcriptomic phylotypic embryonic periods are reported to occur at mid-embryogenic in plants and animals^19–22^, although such chronological staging is especially problematic in plants where embryo-morphogenic outcomes can vary dramatically among taxa^22^. Nonetheless, the question arises as to why older, sequence-conserved genes are preferentially utilized during mid-embryogenesis in both plants and animals? What morphogenetic characteristics embody this mid-embryogenic stage favoring the utilization of ancestral and conserved gene networks?

One concern when answering this question is the temporal delay between gene expression curves and the developmental phenomenology that expression evokes. The maize phylotypic period begins its steep decline in transcriptomic age (TAI) in the proembryo where the inner-and-outer embryonic layers are formed, then culminates during mid-embryogenic Stage 1 when the scutellum and coleoptile are expanding primordia, and the first foliar leaf is just initiating (Fig. 1d). Similarly in *Arabidopsis*, the transcriptomic phylotypic hourglass begins its decent during histological patterning at the globular stage, and peaks during the torpedo stage when the two cotyledon primordia exhibit expansive growth^22^. Moreover, the transcriptomic phylotypic peak of the *P. patens* embryo begins during S2 and peaks during S3, stages comprising expansive growth of the spore capsule from a central meristematic layer and histogenesis of the sporangium (Fig 6). Thus, although no lateral organs are formed in the moss embryo, the transcriptomic phylotypic hourglass in *P. patens* correlates with expansive embryonic growth and histogenesis.

Likewise in *Drosophila* embryogenesis^20,21^, gene age (TAI) peaks during germband elongation^20^, at the onset of embryo segmentation and initiation of the organogenic imaginal discs^49^. Notably, the trend toward the usage of older gene transcripts in the fruit fly begins earlier in embryogenesis, during gastrulation and formation of tissue layers ^20^. A similar pattern is observed in zebrafish (*Danio rerio*), wherein the phylotypic hourglass begins its descent toward older gene transcripts during gastrulation (histogenesis) and then peaks in the subsequent late segmentation/early pharyngula stages, when organ primordia initiate and grow^20^. A later reduction in relative transcriptomic gene age is noted during the zebrafish larval stage, which is accompanied by additional morphogenesis. Taken in summary, the mid-embryogenic, phylotypic period of plants and animals correlates with tissue layer organization and expansive growth of embryonic organs.

Despite their evolutionary distances, we demonstrate that in maize and a moss, as in *Arabidopsis*, fruit flies, and fish, the processes of embryonic tissue-layer histogenesis and primordial growth utilize old and conserved genes, comprising anciently evolved, developmental genetic networks that are proposed to initiate the fundamental body plan^20–22^. Importantly, these phylotypic gene networks are not homologous across phyla and across kingdoms, but rather appear to comprise disparate, old and conserved gene networks that are predicted to generate the variable, phylotypic-staged embryo morphologies seen within animal and plant phylotypes^13, 23^. At the end of this embryonic phylotypic period, plant and animal organs express newer and divergent gene networks in species-specific ways, to generate the vast morphological diversity found in nature.

Our phylotypic analyses of the relatively recently-evolved grass embryo are interpreted to indicate that an evolutionary mechanism for the innovation of morphological novelty exists in plants as well as animals. The maize embryo phylotypic period displays significantly higher TAI values (expression of newer genes) at the Transition and Coleoptilar stages than during Stage 1 (Fig 5b). Notably, no such deviation in the hourglass, phylotypic shape is detected during cotyledon initiation in *Arabidopsis* embryogenesis^22^. These data likely reflect the relatively recent evolution of the bipartite grass cotyledon (the scutellum and coleoptile), which arose long after the cotyledon evolved from an ancestral embryonic leaf 1^3, 4^ and whose evolution involved newer, non-ancestral gene networks. At the same time, the initiating scutellum uses highly-conserved gene networks corresponding to the TDI maximum of maize embryogenesis (Fig. 5c). These data indicate that after the scutellum first appeared, the expression of highly-conserved gene networks during scutellum initiation ensured that this morphologically novel, embryonic organ was retained in subsequent generations. Relatedly speaking, the slight decrease in the use of conserved genes (TDI value) during the Coleoptilar stage may be attributable to morphogenetic differences in the initiating coleoptile ensuring that the bipartite components of the single maize cotyledon (scutellum and coleoptile) are morphologically distinct. We interpret these data to reveal a conserved, testable mechanism involving inverse trends in TAI and TDI values during the innovation of evolutionary novelty in plants and animals.

## Methods

### Plant Materials and growth conditions

B73 inbred lines, the *ns* 1:1 lines (segregate 1/2 wild type NS1+/*ns1 ns2/ns2* plants and 1/2 *ns1/ns1 ns2/ns2* double mutant^50^, and *log7*/+ (obtained from the Maize Genetics COOP Stock Center: UFMu-02863 mu1030680::Mu) were grown in the Gutermann greenhouse facility, Cornell University (Ithaca, NY). Transition staged, coleoptilar staged, Leaf 1-staged, and Leaf 2-staged embryos were collected at 8-9 DAP, 10-11 DAP, 12-13 DAP, 14-15 DAP, respectively. *ns* 1:1 seeds were planted in soil consisting of a 1:1 mixture of Turface and LM111 in Percival A100 growth chamber at Weill Hall, Cornell University (Ithaca, NY) with the set condition: day 29.4 °C/16 hours, night 23.9 °C/8 hours, humidity 50%. The seedlings were collected on the 7^th^ day after planting for coleoptile comparisons between WT and *ns1;2* mutants.

### Single-cell transcriptomic analysis

Two batches of B73 Leaf1 staged embryos were collected in summer 2021 and winter 2022. For each batch, 200 embryos were dissected from developing seeds for protoplast isolation as previously described^51^. Protoplast (1000-2000 cells per µL) were loaded into the 10X Genomics Chromium platform to generate a Gel Bead-In-Emulsion (GEM). The GEM-captured RNA was reverse transcribed and amplified to make 3′ cDNA libraries, followed by the high throughput RNA sequencing on the Illumina NextSeq 2000 sequencer at the Cornell University Biotechnology Resource Center.

The resulting raw base call (BCL) files were demultiplexed to generate FASTQ files via CellRanger mkfastq v6.0 (10x Genomics, Pleasanton, CA). FASTQ reads were trimmed, aligned (to B73 reference genome V3^52^) and assigned to cell barcodes to generate expression count matrices via CellRanger count with default settings. The expression count matrices were then analyzed using Seurat v4^53–55^. In detail, the matrices were filtered with the setting (number of genes > 1500, UMI > 6000) to remove low quality cells. The filtered matrices were normalized and log transformed via SCTransform function. The normalized matrices were merged, and batch effects were removed via R package Harmony v1.2.0^56^. The cell-cycle associated genes were scored and regressed via CellCycleScoring and ScaleData function. Matrix dimension was performed using RunPCA. The cell clusters and the Uniform Manifold Approximation and Projection (UMAP) plot were generated with the setting (dim = 1:40, resolution = 1, n. neighbors = 40, min. dist = 0.01, spread = 3). The differential expressed genes and cell cluster markers were identified via FindAllMarkers function with the log2 fold change threshold = 0.2.

### Spatial transcriptomic analysis

For use in spatial transcriptomics, B73 Leaf1 staged embryos were processed for either paraffin-embedding and microtome sectioning, or cryoembedding and cryosectioning. For paraffin-embedding, two embryos were fixed in Farmer’s Fixative (Ethanol:Acetic acid = 3:1), followed by dehydration in an ethanol concentration gradient (75%, 85%, 95%, 100%) and clearing with Ethanol:Xylene gradient of 3:1, 1:1, 1:3, and 100% Xylene at room temperature. Cleared embryos were embedded in Paraplast Plus® (Leica Biosystems, Deer Park, IL). The embedded blocks were microtome-sectioned at 10 um at room temperature; sections were subsequently mounted onto 10x Visium^TM^ Spatial Gene Expression Slides (10x Genomics, Pleasanton, CA). Slides were deparaffinized with two changes of 100% Xylene and 95% Ethanol for 10 min and 2 min, respectively. For cryoembedding, two kernels were dissected longitudinally to remove left and right margin of the kernel; the central region containing the embryo was embedded in Tissue-Tek® Optimal Cutting Temperature (O.C.T., Sakura Finetek, Torrance, CA), and snap-frozen in the dry ice. Frozen tissue blocks were cryosectioned at10 µm, and the sections were mounted on 10x Visium^TM^ Spatial Gene Expression Slide (10x Genomics, Pleasanton, CA).

The 10x Visium^TM^ Spatial Gene Expression Slides were processed for spatial library construction following the manufacturer instructions. Briefly, the sections were baked at 37 °C for 1 min followed by re-fixation in 100% methanol at –20 °C for 30 min. Subsequently, the sections were stained with Toluidine Blue for histological imaging. Tissue permeabilization was performed at 37 °C for 6 min to release RNA from the fixed tissue, followed by first strand cDNA synthesis. Subsequently, second strand cDNA was synthesized and amplified for library construction and sequencing. High-throughput RNA sequencing was conducted on an Illumina NextSeq 2000 sequencer at the Cornell University Biotechnology Resource Center.

Raw reads were demultiplexed, mapped to the Toluidine Blue stained images, and aligned to the B73 reference genome V3^52^ to generate spatial expression count matrices using SpaceRanger v2.0 (10x Genomics, Pleasanton, CA). The matrices were analyzed using Seurat v4^53–55^. In detail, the matrices were normalized and log transformed via SCTransform function. The normalized matrices were merged followed by dimensionality reduction. Cell clusters and the UMAP were generated with the following settings (dim = 1:40; resolution = 1.5; neighbors = 10; minimum distance =0.01). The spatial data were visualized with DimPlot and SpatialPlot; differential expressed genes and cell cluster markers were identified via FindAllMarkers function with the log2 fold change threshold = 0.1.

### Laser-microdissection RNAseq analysis

Sixty maize Leaf1 staged kernels in two replicates were dissected longitudinally to remove the left and right margins of the kernels. This embryo-containing central region was fixed in Farmer’s Fixative (Ethanol:Acetic acid=3:1). After fixation, the samples were dehydrated, cleared, embedded and sectioned as described previously^57, 58^. Tissue sections were mounted on PEN membrane slides (Leica Microsystems, Deer Park, IL) pre-treated at 180 °C for 4 hours followed by UV irradiation at 254 nm for 30 minutes. The sections were deparaffinized with two changes of 100% Xylene and 100% Ethanol for 10 min and 2 min, respectively. Laser microdissection was performed using a PALM Laser Microbeam System (Zeiss Group, Oberkochen, Germany). Adaxial scutellum, SAM, leaf, as well as upper and lower coleoptiles were microdissected (total area for each captured organ is over 0.3 mm^2^) and RNA extraction was performed using the Arcturus^TM^ PicoPure^TM^ RNA Isolation Kit (Thermo Fisher Scientific, Waltham, MA). The extracted RNA was amplified using Arcturus^TM^ RiboAmp^TM^ HS PLUS RNA Amplification Kit (Thermo Fisher Scientific, Waltham, MA). RNA quality control, library construction and sequencing were performed at the Cornell University Biotechnology Resource Center. The raw reads quality control was conducted via FastQC (https://www.bioinformatics.babraham.ac.uk/projects/fastqc/). Raw reads were mapped to the. B73 reference genome V3^3^ by HISAT2 (v2.1.0)^59, 60^. Read counts and the normalized number of transcripts (transcript per million reads, TPM) were calculated as previously described^61^. DEseq2 (v1.26.0)^62^ was used to identify differentially expressed genes (DEGs) with the cutoff criteria of log2 fold change >1 or <1 with the false discovery rate (FDR) < 0.05.

### Multiplex *in situ* hybridization

Transition staged, coleoptilar staged and Leaf1 staged kernels (two replicate for each stage) were dissected longitudinally to remove the left and right kernel margins. The central region containing the embryo was fixed in 4% paraformaldehyde (PFA); fixed samples were dehydrated, cleared, embedded and sectioned as described previously^63, 64^. Sections were mounted on 10x Genomics Xenium^TM^ slides, deparaffinized and de-crosslinked following the manufacturer’s instructions. The Xenium custom gene expression panel containing probes of 99 maize genes were applied on each section to perform hybridization, ligation and rolling circle amplification, followed by cycles of fluorescent probe hybridization, imaging and decoding. The data was visualized by Xenium Explorer v1.3. The Xenium^TM^ multiplex *in situ* hybridization assays were conducted at Memorial Sloan Kettering Cancer Center (New York, NY).

### Histology and immunohistolocalizations

Developing kernels from the proembryo stage, transition stage, coleoptilar stage, and the Stage 1 were fixed in FAA, dehydrated, cleared, embedded in paraplast, sectioned and stained as described previously^64^. Immunohistolocalizations were performed using an antibody to maize Class I KNOX proteins, KNOTTED1 and ROUGH SHEATH1, as described^65^.

### WGCNA and hdWGCNA

For WGCNA to pursue to initiation network, log2-transformed TPM data from LM-RNAseq (SAM data was excluded) were filtered following these criteria: row mean > 0.2 and coefficient of variation > 0.4. The general pipeline was performed as per R package instructions^40^ with minor modifications as follows. The network type was set to “signed”. The soft power, the minimum module size, and the merging threshold were set to 13, 20 and 0.3, respectively. For the analysis to pursue transcriptomic relationship between organs, the LM-RNAseq data in this study was merged with the LM-RNAseq data of the 14-day seedling P0 and P1 data^42^. The data was normalized and batched effect removed as described, and filtered following the same criteria as the WGCNA for the initiation network. The network type was set to “signed”. The soft power, the minimum module size, and the merging threshold were set to 16, 20 and 0.4, respectively.

For hdWGCNA, the single-cell RNAseq data was used to perform the analysis. The data was SCTransformed and normalized. The general pipeline was conducted as the R package instructions^43^ with the following minor modifications. The network type was set to “signed”, and the soft power and minimum module size were set to 10 and 30, respectively. Hypergeometric test was performed to analyze the correlation between the hdWGCNA modules and the single-cell clusters following the previously described procedures^66^.

### Statistical analysis

Hypergeometric tests were performed to analyze the correlation between single-cell clusters and spatial transcriptomic clusters, as well as between the single-cell clusters and hdWGCNA modules. We followed previously algorithms^66^ to statistically evaluate if the two groups of data were significantly correlated by calculating the number of overlapping gene relative to the total number of protein-coding genes in the maize B73 reference genome V3^52^. The p values were transformed by –log10p, and visualized as heatmap built by the R package gplots (https://github.com/cran/gplots).

LM-RNAseq read count data were normalized to fit the generalized linear model (GLM), and Empirical Bater shrinkage estimation was used for calculating dispersions, expression fold changes and adjust p values. In addition, Pearson correlation test was used to calculate the correlation between the coexpression modules and the corresponding clusters (microdissected organs or putative initiation network) in WGCNA, and between the maize and Arabidopsis embryogenesis genes.

To statistically compare the expression of the initiation network between the transition, coleoptilar and the stage1, the TPM data was log2 transformed [log2(TPM+1)] and compared using Tukey-Kramer HSD test by JMP Pro 17 (SAS Institute, Cary, NC) with default settings.

For the correlation test, the transition, coleoptilar and the stage1 TPM data was double log2 transformed, i.e. log2[log2(TPM+1)+1], to fix the data skewness caused by the outliers. The Arabidopsis TPM data was log2 transformed. The Pearson correlation was performed in R v4.3.2 with the significance cutoff value p<0.05.

### Calculating TAI and TDI for phylotypic period

The procedures of constructing a phylostratigraphic map, calculating TAI and TDI, and statistical tests, have been presented previously^20,22,45,67^. For each of the thirteen phylostrata for maize (Fig. 5a), the amino acid sequences of all 350 species (Supplementary Dataset 15) with completely sequenced genomes were extracted from NCBI, Ensembl genomes, Treegene (https://treegenesdb.org), Fernbase (https://fernbase.org), or Maizegdb (https://maizegdb.org). An equivalent strategy was applied for the five *P. patens* phylostrata f(Fig. 6b), encompassing 174 species (Supplementary Dataset 17). Each of the 63,031 amino acid sequences in *Zea mays* ssp. *mays* (v3) and 24, 002 sequences in *P. patens* (v1.6) with minimum length of 30 amino acids was analyzed by BLASTp (BLAST version 2.15.0) with an E-value cut-off of 10^-5^. Each such gene of *Zea mays* ssp. *mays* was assigned to the phylogenetically most distant (oldest) phylostratum containing at least one species with at least one blast hit. In circumstances where no blast hit was identified in older phylostrata, the *Zea mays* ssp. *mays* gene was assigned to phylostratum 13 (PS13; Fig. 5a). In this way, each gene of *P. patens* was likewise assigned to one *P. patens* phylostratum, from PS1 to PS5 (Fig 6b).

To evaluate gene sequence divergence, the *K_a_/K_s_* ratio was calculated between maize and its sister species *Zea diploperennis* or *Zea Mexicana*, between *Physcomitrium patens* and the closely-related species *Physcomitrellopsis africana*^68^. *K_a_* and *K_s_*refer to the number of non-synonymous and synonymous substitutions, respectively, between each orthologous pair. Orthologous gene pairs were determined with the method of best hits using BLASTp. Amino acid sequence alignments of each pair generated with MAFFT^69^ (L-INS-i option) were used for codon alignments generated with PAL2NAL^70^ to compute sequence divergence levels (Ka/Ks) with KaKs Calculator (version 3.0)^71^. Gene pairs with K_a_<0.5, K_s_<5 and K_a_/K_s_ ratios < 2 were retained.

Transcriptomic data for maize embryonic developmental stages were collected from four data sources 1) LM-RNAseq data of proembryo stage^28^, 2) organ initiation network (this study), 3) RNAseq data of Stage 2 embryo^72^ and 4) LM-RNAseq data of 14-day after germination seedling P0 and P1 leaves^42^. The datasets were processed with batch effect removal and normalization as described above. The TAI and the TDI are weighted means of evolutionary age and sequence divergence, respectively, and defined previously^20, 22^. The transcriptome age index TAI_s_ and the transcriptome divergence index TDI_s_ of developmental stage s (s = proembryo, transition, coleoptilar, stage 1 leaf, stage 1 scutellum, stage 1 coleoptile, seedling leaves P0 and seedling leaves P1) are the weighted mean of the evolutionary age (phylostratum) weighted by the expression level of each gene at developmental stage s, and the weighted mean of the Ka/Ks ratio of each gene weighted by the expression level of each gene at developmental stage *s*, respectively. Transcriptomic data for embryo/sporophyte stages in *P. patens* were from Ortiz-Ramirez et al., (2016)^46^. Low/high PS values correspond to evolutionarily old/young genes, so low/high TAI values correspond to evolutionarily old/young transcriptomes. Low/high Ka/Ks ratios correspond to conserved/divergent genes, so low/ high TDI values correspond to conserved/divergent transcriptomes. For gene models with multiple predicted peptides, TAI was calculated with the most conserved phylostratum assigned to that locus and TDI was calculated with the conserved codon divergence strata assigned to that locus.

To determine the statistical significance of the TAI and TDI profiles, the flat line test^22^, reductive hourglass test and reductive early conservation test were performed via myTAI package (v1.0.1)^67^. The flat line test is a permutation test based on the variance of the TAI or TDI values of a given TAI or TDI profile. For the reductive hourglass test in maize, the embryonic stages were partitioned into three modules, early (proembryo stage), mid (transition stage, coleoptilar stage, Stage 1 leaf, Stage 1 scutellum, and Stage 1 coleoptile), and late (Stage 2 and seedling leaves), and a permutation test was performed for the minimum differences of the mean TAI or TDI values between early and mid-modules, and between late and mid modules. For the reductive early conservation test in maize, a permutation test was performed for the minimum differences of the mean TAI or TDI values between mid and early modules, and between late and early modules. For the reductive hourglass test and reductive early conservation tests on *P. patens*, the three modules comprised the archegonium and S1 stage (early), S2 and S3 (middle), and SM (late). Detailed procedures are described^67^.

## Data availability

The single-cell RNAseq, Visium^TM^ spatial transcriptomics, and LM-RNAseq data were deposited in the NCBI Sequence Read Archive (SRA) with the BioProject ID: PRJNA1045558. The Xenium^TM^ multiplex *in situ* hybridization data was deposited in Science Data Bank with the link: https://www.scidb.cn/s/F7ZbEz, which could be visualized by Xenium Explorer v1.3.

## Code availability

The code for TAI and TDI calculations was uploaded to https://gitlab.com/rz444/maizephylper.git.

## Supplementary figure legends

**Supplementary Fig. 1.**
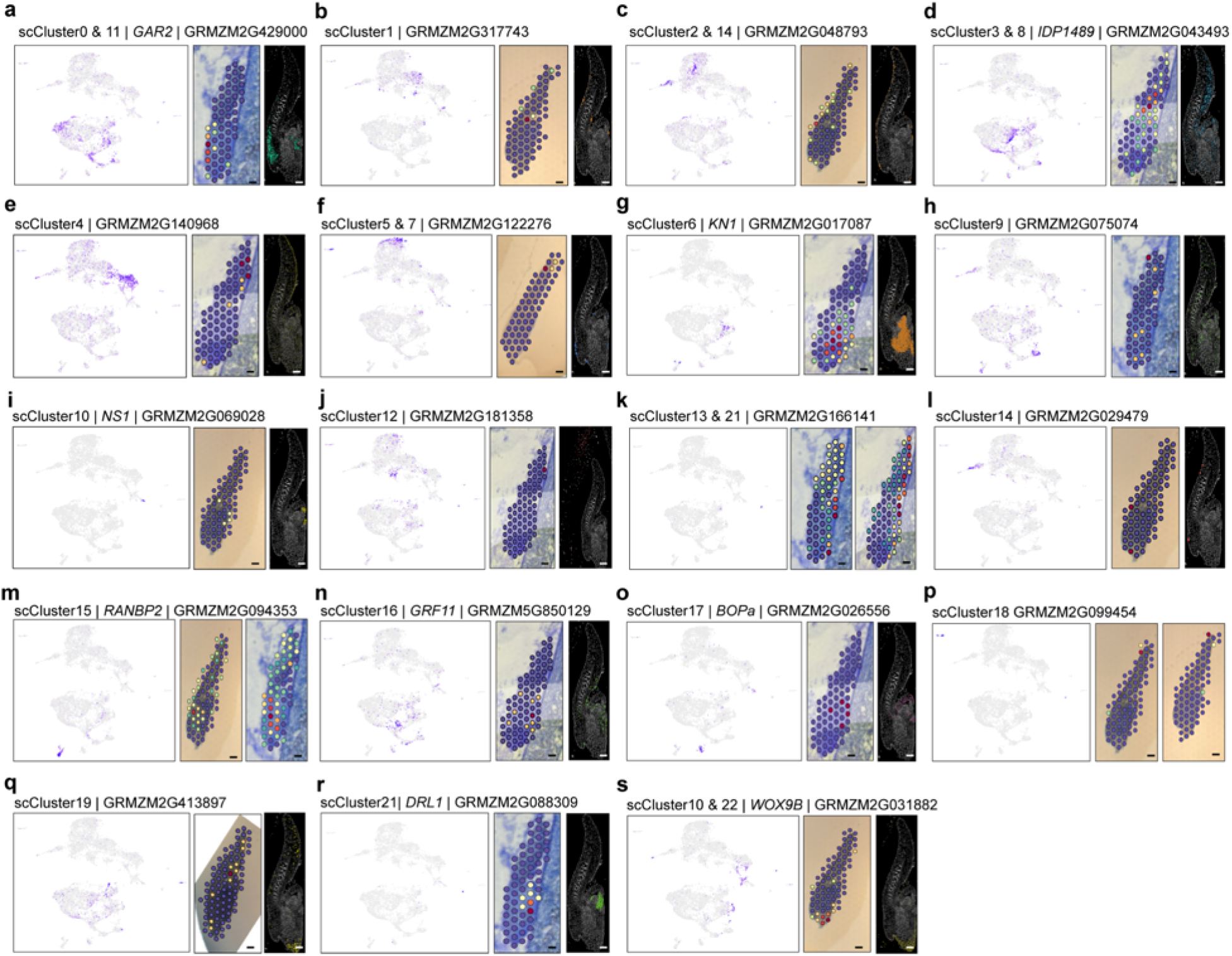
Verification of single-cell cluster marker genes. Left panel: single-cell cluster marker expression projected as a UMAP. Purple dots mark cells with corresponding gene-specific expression. Middle panel: verification of single-cell cluster marker genes by Visium^TM^. Red to green spots mark the locations of gene expression with high to low level, respectively. Right panel: verification of single-cell cluster marker genes by multiplexed RNA-targeting *in situ* hybridization [data not available for *GRMZM2G166141* (**k**), *GRMZM2G094353* (**m**) and *GRMZM2G099454* (**p**)]. Scale bar = 100 μm.

**Supplementary Fig. 2.**
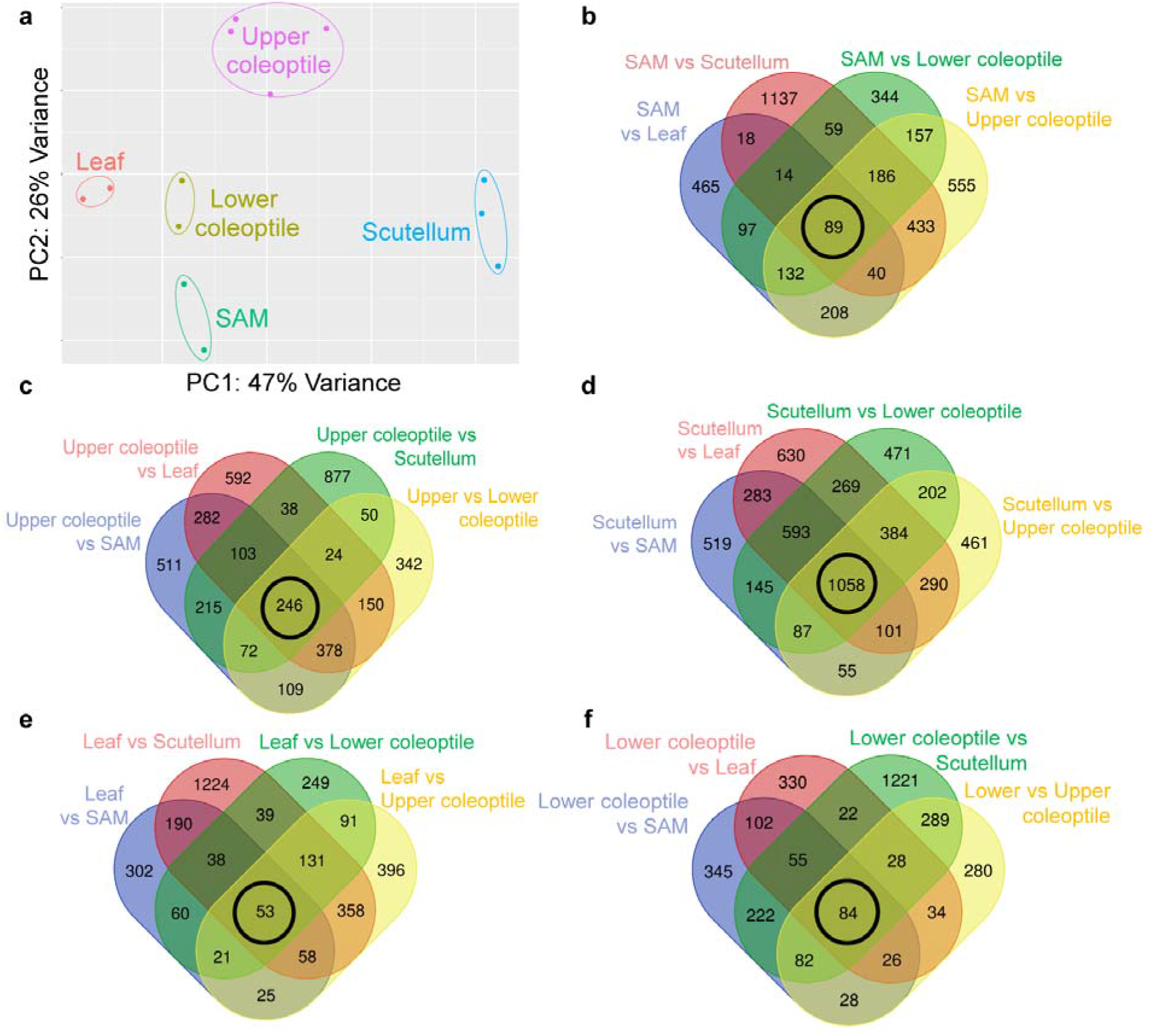
Differential expression between laser microdissected RNAseq samples. **a**. Principal component analysis of laser-microdissected RNAseq samples. Each dot represents a laser-microdissected embryonic shoot sample. **b-f**. Pairwise comparison of the gene expression between laser-microdissected RNAseq samples. The number of overlapping, differential-expressed genes (DEGs) is showed in the Venn diagram. The circled numbers represent the genes preferentially expressed in the SAM (**b**), upper coleoptile (**c**), scutellum (**d**), leaf (**e**) and lower coleoptile (**f**).

**Supplementary Fig. 3.**
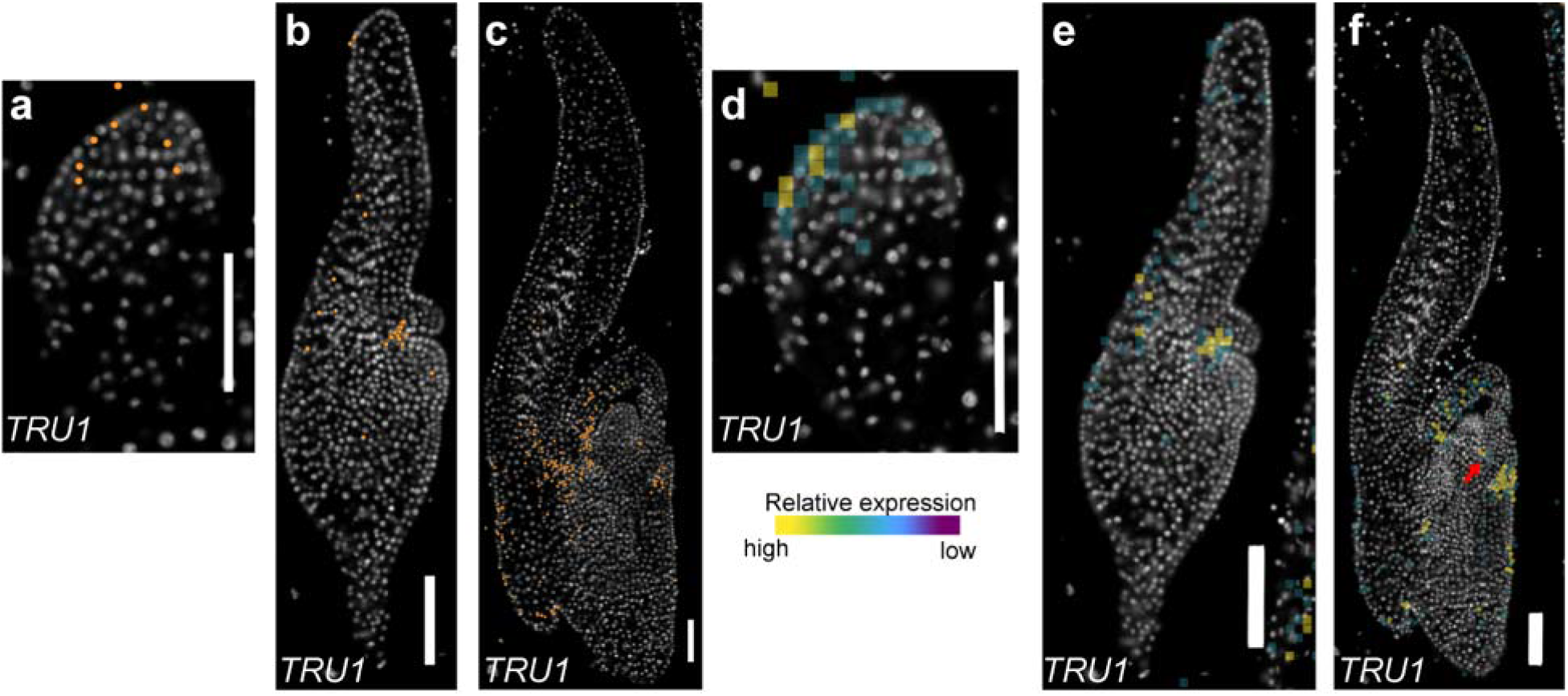
Multiplex *in situ* hybridization of *TRU1*. Spatio-temporal expression of *TRU1* displayed as a signal point map (**a-c**) and as a transcript abundance correlated signal density map (**d-f**) at the transition stage (**a and d**), coleoptilar stage (**b and e**) and Stage 1 (**c and f**). The red arrow in (f) marks the expression of *TRU1* at the emerging leaf sheath of leaf 1. Scale bar = 100 μm.

**Supplementary Fig. 4.**
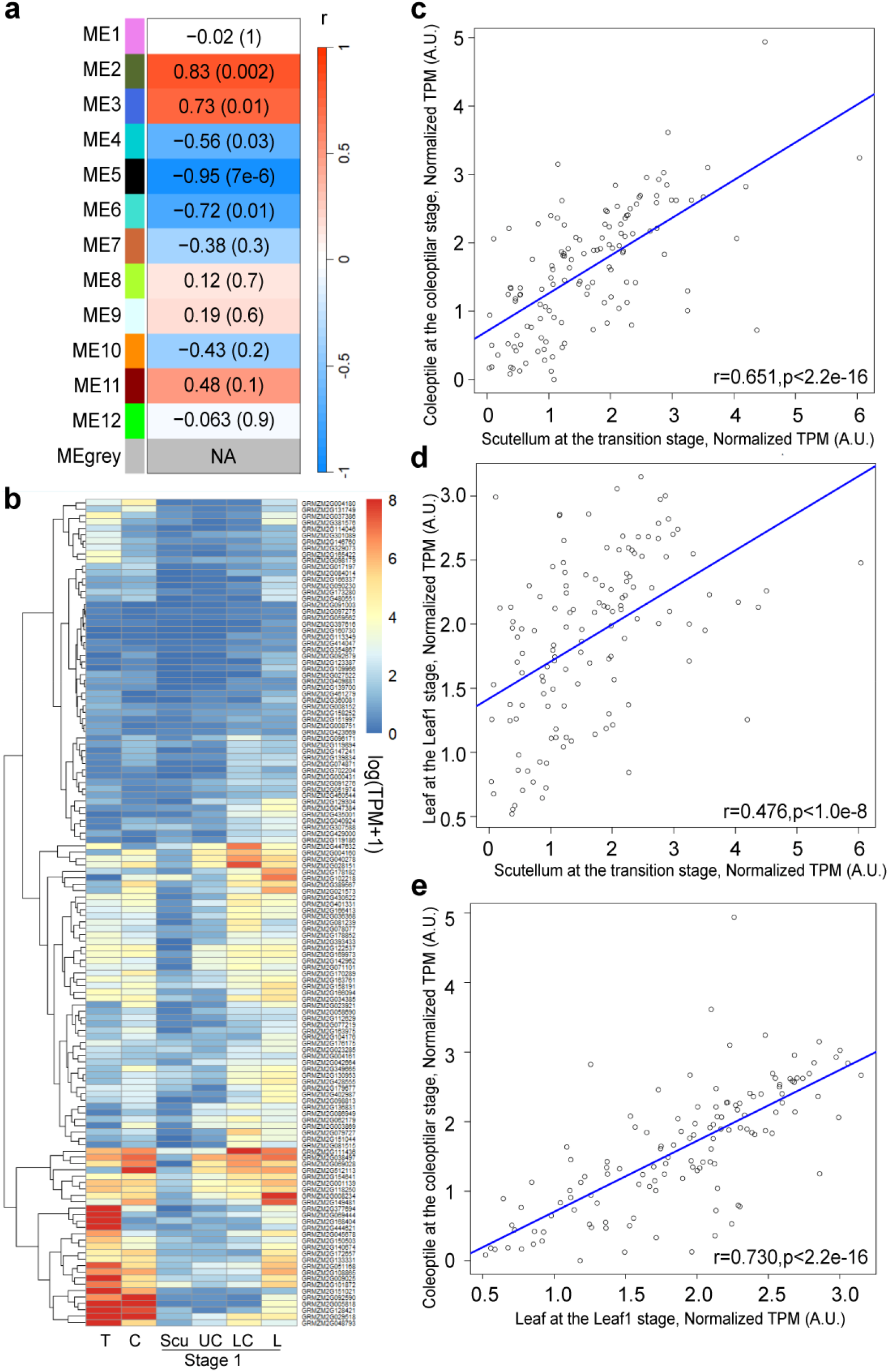
Expression of the embryo organ initiation network. **a**. WGCNA identifies modules with overall highest expression at Stage 1 leaf (L), less in the lower coleoptile (LC), and steadily decreased expression in the upper coleoptile (UC), scutellum (S). Heatmap shows the correlation coefficients (r) and the corresponding p values (parentheses) of modules ME1-12. **b**. Expression of the 130 genes comprising the embryonic-organ-initiation network at the transition stage (T), coleoptilar stage (C) and the Stage 1 scutellum (Scu), upper coleoptile (UC), lower coleoptile (LC) and leaf 1 (L). **c-e**. Correlations of the expression of these 130 genes during initiation of the scutellum, coleoptile, and leaf 1.

**Supplementary Fig. 5.**
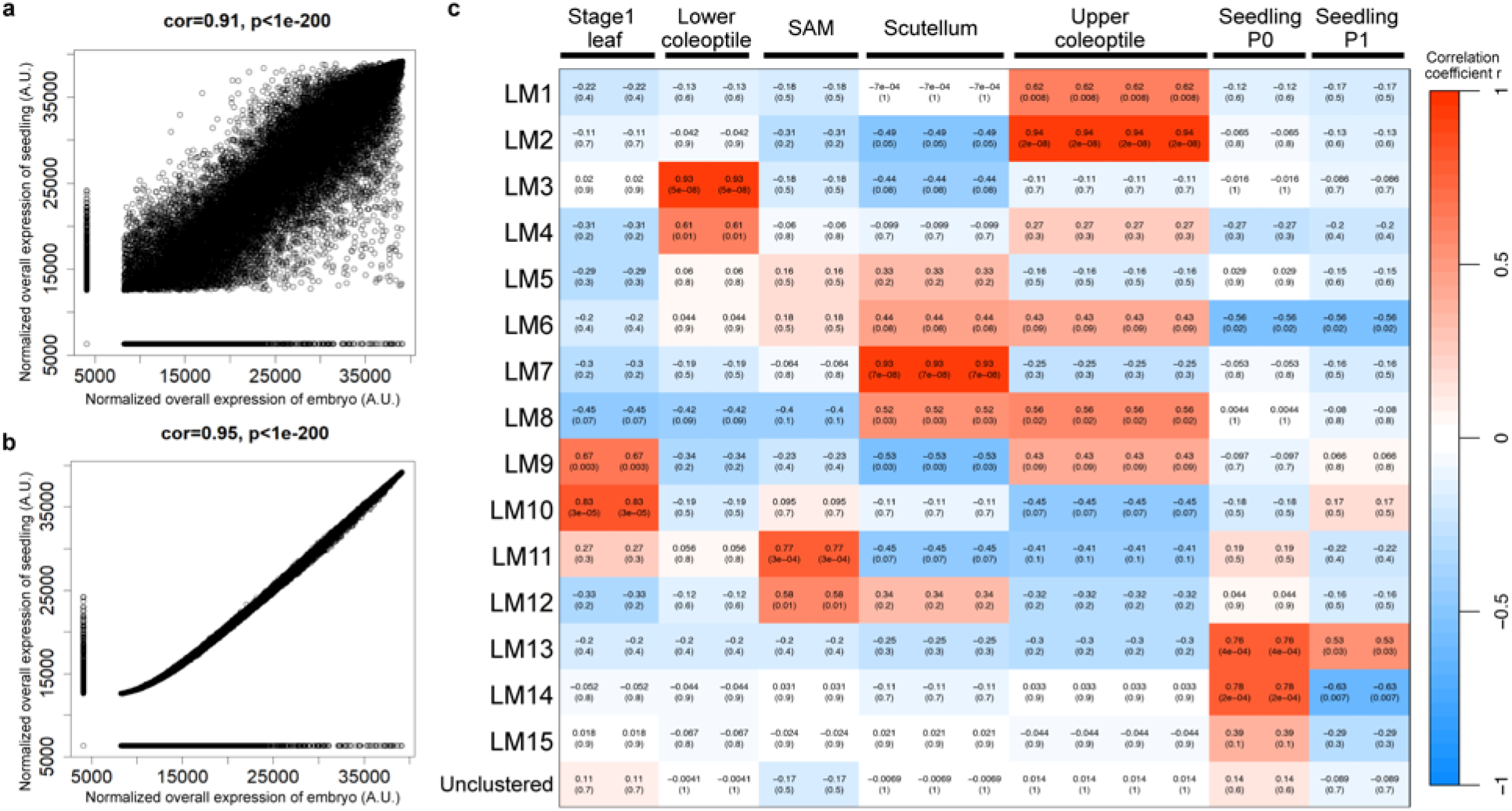
WGCNA Module-organ correlation. **a-b**. Correlations of transcriptomic datasets from embryonic organs and leaves initiated post-germination before and after batch effect removal. Correlations of the normalized overall gene expression between the post-germination seedling leaves and embryonic stages before (**a**) and after (**b**) batch effect removal. **c**. Heatmap highlights correlations between co-expression modules (y-axis) and corresponding organs (x-axis). Colors are graded based on correlation coefficients (r) and p values (in parentheses).

**Supplementary Fig. 6.**
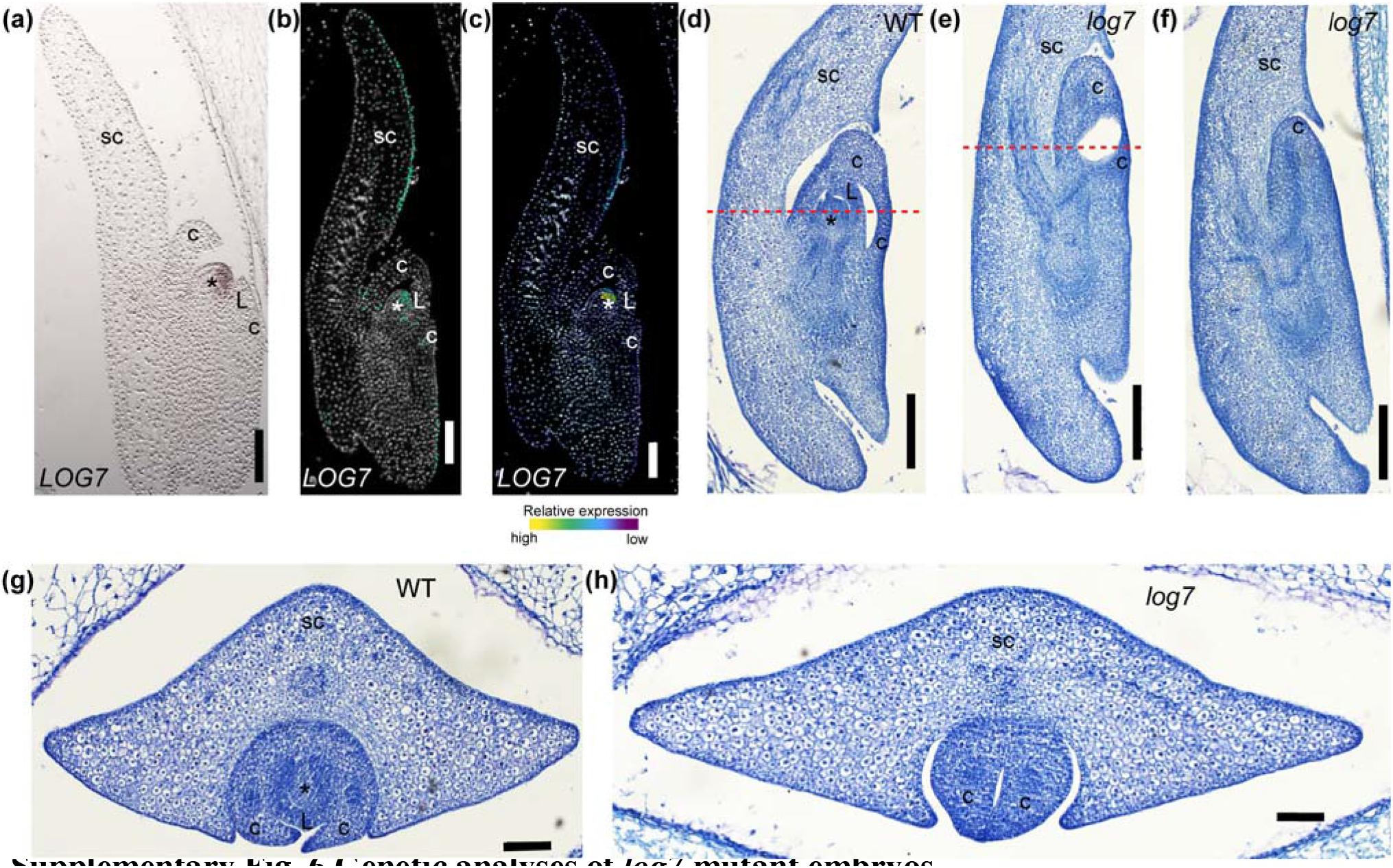
Genetic analyses of *log7* mutant embryos. Expression and genetic analysis of *LOG7***. a.** Conventional *in situ* hybridization of *LOG7* transcript accumulation in a longitudinal section of the Stage 1 embryo. Multiplexed RNA-targeting point map (**b**) and density map (**c**) of *LOG7* accumulation in longitudinal sections of Stage 1 embryos. Scale bar = 100 μm. **d.** Longitudinal median section of a WT Stage 2 embryo. The red dashed line marks the transverse section plane depicted in (**g**). **e-f.** Longitudinal median sections of the *log7* mutant Stage 2 embryos. The red dashed line marks the transverse section plane depicted in (**h**). Longitudinal off-median section of the *log7* mutant Stage 2 embryo. All scale bars = 100 μm. Asterisks (*) mark the SAM; sc, scutellum; c, coleoptile; L, leaf.

**Supplementary Fig. 7.**
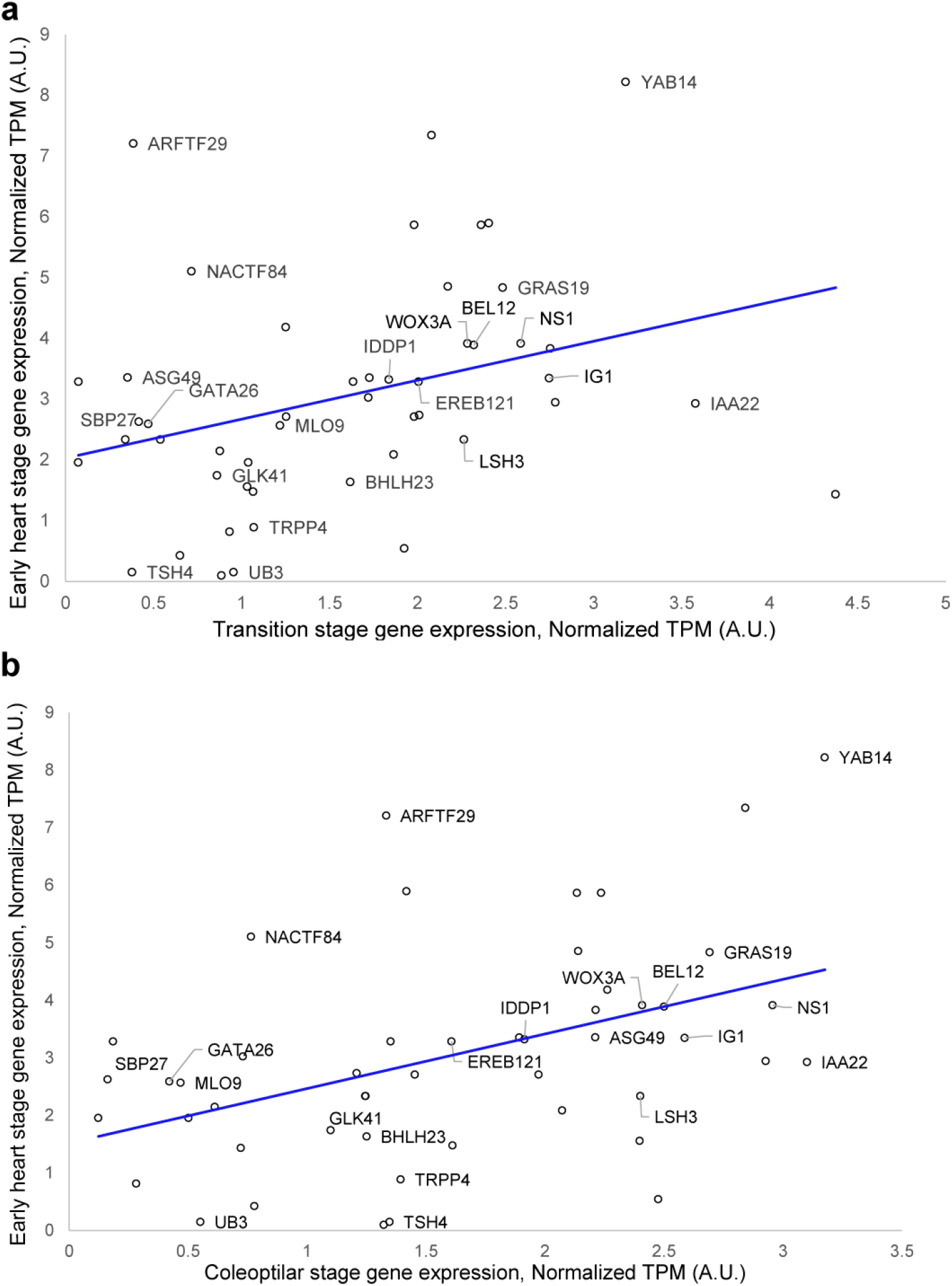
Correlation between the maize initiation network genes and the Arabidopsis orthologs. The correlation of expression between the maize initiation network genes at transition (**a**) and coleoptilar (**b**) stage, and the Arabidopsis orthologs at early heart stage.

**Supplementary Fig. 8.**
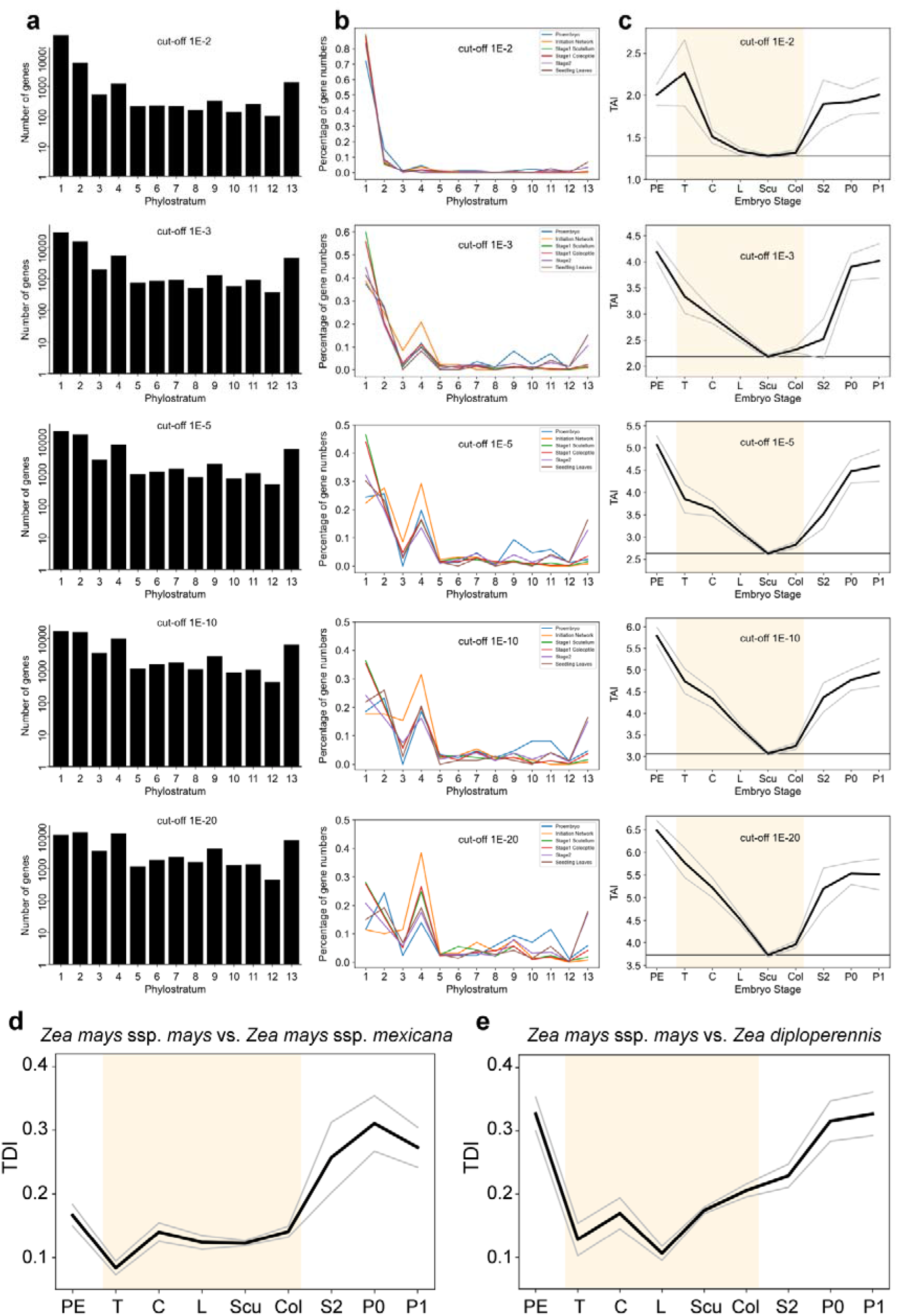
TAI and TDI for embryo stages under different BLASTp E-value cutoffs. **a-c**. Distribution of all maize genes and genes in each embryo stage across phylostrata and TAI profiles for different E-value cut-offs in BLASTp homology searches (ranging from 1E-2 to 1E-20). This analysis investigates whether and how different E-values affect the hourglass pattern. **a**. Variation of the E-value cut-off for the BLASTp analysis demonstrates minimal effects on the distribution of maize genes among the phylostrata, indicating an overall pattern independent of the E-values cut-offs. The y-axis is presented on a logarithmic scale. **b**. Similarly, the variation of the E-value cut-off for the BLASTp analysis does not severely alter the distribution of maize genes in each embryo stage among the phylostrata. **c**. The hourglass pattern of the TAI profile remains consistent across various E-value cut-offs. **d-e**. Transcriptome divergence indices based on two comparisons of genomes from three relatives of *Zea mays*: *Zea mays* ssp. Mays vs. *Zea mays* ssp. mexicana (**d**); *Zea mays* ssp. mays vs. *Zea diploperennis* (**e**). The overall patterns of TDI profiles are significant, as measured by the flat line test (*P*_TDI-Mexicana_ = 1.07×10^-3^, *P*_TDI-diploperennis_ = 4.80×10^-4^), reductive hourglass test (*P*_TDI-Mexicana_ = 0.027, *P*_TDI-diploperennis_ = 1.53×10^-3^), and reductive early conservation test (*P*_TDI-Mexicana_ = 0.68, *P*_TDI-diploperennis_ = 0.998).

## Supplementary information

Supplementary Table 1. Single-cell RNAseq cluster marker genes

Supplementary Table 2. Identity of single-cell clusters

Supplementary Table 3. Spatial transcriptomic cluster markers

Supplementary Table 4. Normalized gene expression (TPM) of LM-RNAseq

Supplementary Table 5. Normalized gene expression (log2 (TPM+1)) of tissue specific classic genes

Supplementary Table 6. Gene expression of selected leaf genes in the transition, coleoptilar and Leaf 1 stage in the LM-RNAseq data

Supplementary Table 7. Xenium^TM^ multiplex in situ hybridization probes and result

Supplementary Table 8. WGCNA: modules with gene expression pattern: Scu < UpCol < LoCol < Leaf

Supplementary Table 9. Putative initiation network and the log2-transformed TPM in the lateral organs of the transition, coleoptilar, Stage 1 (L1) and 14-day after germination seedlings

Supplementary Table 10. WGCNA: Cell-type specific modules

Supplementary Table 11. hdWGCNA module table

Supplementary Table 12. Maize initiation network vs Arabidopsis orthologs

Supplementary Table 13. Genomes used for phylogeny reconstruction and phylostratigraphic map of maize

Supplementary Table 14. Phylostrata of maize genes

Supplementary Table 15. The gene numbers of each phylostratum in maize embryo stages

Supplementary Table 16. Gene expression for TAI and TDI calculation in maize

Supplementary Table 17. *Ka/Ks* ratio of maize genes expressed at the embryogenesis and 14-day seedling

Supplementary Table 18. Genomes used for phylogeny reconstruction and phylostratigraphic map of *Physcomitrium patens*

Supplementary Table 19. Phylostrata of *Physcomitrium patens* genes

Supplementary Table 20. Ka/Ks of *Physcomitrium patens* genes

## Acknowledgements

We thank L. Evans and R. Ragas for helpful discussions of the data and comments on the manuscript. Special thanks to Cornell Genomics Facility for performing single-cell and Visium™ sequencing, to the Cornell Transcriptional Regulation and Expression Facility for RNA sequencing, and the Cornell Guterman Bioclimatic Labs for our plants. Thanks to the Integrated Genomics Operation (IGO) of Memorial Sloan Kettering Cancer Center for performing Xenium assays. H. Wu was supported by a grant from the National Science Foundation (IOS-2016021). R. Zhang is supported by another grant from the National Science Foundation (IOS-2210259).

## Author contributions

Author Contributions M.J.S., H.W., R.Z., and K.J.N. conceived the study. H.W., M.J.S., and R.Z. generated and analyzed the data. M.J.S, H.W., R.Z. and K.J.N. interpreted the results, and M.J.S, H.W., R.Z., and K.J.N. wrote and edited the manuscript.

## Ethics declaration

Competing interests

The authors declare no competing interests.

## Corresponding author

Michael J. Scanlon

Hao Wu

